# SARS-CoV-2 Infects Human Engineered Heart Tissues and Models COVID-19 Myocarditis

**DOI:** 10.1101/2020.11.04.364315

**Authors:** Adam L. Bailey, Oleksandr Dmytrenko, Lina Greenberg, Andrea L. Bredemeyer, Pan Ma, Jing Liu, Vinay Penna, Lulu Lai, Emma S. Winkler, Sanja Sviben, Erin Brooks, Ajith P. Nair, Kent A. Heck, Aniket S. Rali, Leo Simpson, Mehrdad Saririan, Dan Hobohm, W. Tom Stump, James A. Fitzpatrick, Xuping Xie, Pei-Yong Shi, J. Travis Hinson, Weng-Tein Gi, Constanze Schmidt, Florian Leuschner, Chieh-Yu Lin, Michael S. Diamond, Michael J. Greenberg, Kory J. Lavine

**Author notes:** Co-first authors (Adam L. Bailey, Oleksandr Dmytrenko, Lina Greenberg). Co-senior authors (Chieh-Yu Lin, Michael S. Diamond, Michael J. Greenberg, Kory J. Lavine). **Corresponding authors information:** Kory J. Lavine, MD, PhD, 660 South Euclid Campus Box 8086, St. Louis, MO, 63110, Michael J. Greenberg, PhD, 660 South Euclid Campus Box 8231, St. Louis, MO, 63110, Michael S. Diamond, MD, PhD, 660 South Euclid Campus Box 8051.

## Abstract

Epidemiological studies of the COVID-19 pandemic have revealed evidence of cardiac involvement and documented that myocardial injury and myocarditis are predictors of poor outcomes. Nonetheless, little is understood regarding SARS-CoV-2 tropism within the heart and whether cardiac complications result directly from myocardial infection. Here, we develop a human engineered heart tissue model and demonstrate that SARS-CoV-2 selectively infects cardiomyocytes. Viral infection is dependent on expression of angiotensin-I converting enzyme 2 (ACE2) and endosomal cysteine proteases, suggesting an endosomal mechanism of cell entry. After infection with SARS-CoV-2, engineered tissues display typical features of myocarditis, including cardiomyocyte cell death, impaired cardiac contractility, and innate immune cell activation. Consistent with these findings, autopsy tissue obtained from individuals with COVID-19 myocarditis demonstrated cardiomyocyte infection, cell death, and macrophage-predominate immune cell infiltrate. These findings establish human cardiomyocyte tropism for SARS-CoV-2 and provide an experimental platform for interrogating and mitigating cardiac complications of COVID-19.

## Introduction

Severe Acute Respiratory Syndrome Coronavirus 2 (SARS-CoV-2) is the cause of the ongoing Coronavirus Disease-2019 (COVID-19) pandemic. Since its emergence in Wuhan, China in late 2019, SARS-CoV-2 has infected millions of people worldwide, overwhelmed the capacity of healthcare systems, and continues to result in unacceptably high mortality rates. Although most severe cases of COVID-19 are characterized by respiratory distress, the disease has a highly variable clinical course with myriad manifestations. Epidemiological studies have identified pre-existing cardiovascular disease as a strong risk factor for the development of severe COVID-19 and mortality (reviewed in^1^). Cardiovascular manifestations of COVID-19 including elevated troponin, reduced left ventricular systolic function, and arrhythmias are consistent with cardiomyocyte injury. Cardiac complications occur in 20-44% of hospitalized patients, and constitute an independent risk factor for COVID-19 mortality^1–4^. Cardiac MRI studies have suggested that persistent myocardial injury may be more common than appreciated and occur in less severe forms of COVID-19^5–7^. The mechanistic basis by which SARS-CoV-2 results in cardiac dysfunction remains obscure. It remains unknown whether these effects are a result of direct viral infection of cardiac cells or a systemic inflammatory response to extracardiac infection^8^. If direct infection does contribute to COVID-19-associated cardiac injury, defining the cell-types susceptible to infection will be important for understanding COVID-19 cardiac pathogenesis and devising effective treatment strategies.

It has been challenging to study the cardiac manifestations of COVID-19. Obtaining cardiac tissue from critically ill patients with suspected COVID-19 myocarditis poses unique challenges. Furthermore, there are few animal models to study cardiovascular complications observed in SARS-CoV-2 infected humans^9^. The most commonly used laboratory animal model, the mouse, is not susceptible to SARS-CoV-2 infection due to poor affinity of the viral spike protein for the murine ACE2 receptor^10,11^, and systems that enable expression of human ACE2 from cells in the murine cardiovascular system are not yet widely available. Therefore, there is a critical need to develop robust model systems that enable the investigation of the cardiovascular complications of COVID-19.

Human engineered heart tissues (EHTs) provide unique advantages as model systems for studying COVID-19 cardiac pathology. EHTs are self-assembled using defined cellular and extracellular matrix compositions. EHTs generate contractile force, display electrical coupling, and have cellular organization that mimics myocardial tissue (reviewed in^12,13^). Human EHTs can be formed from human pluripotent stem cell (hPSC)-derived cardiomyocytes, where the three-dimensional environment of the EHT promotes the maturation of these cells^14^.

Here, we devised an EHT model of COVID-19 myocarditis and tested the hypothesis that SARS-CoV-2 promotes cardiac pathology by infecting cardiomyocytes and activating local immune responses. We demonstrate that SARS-CoV-2 selectively infects and replicates within hPSC-derived cardiomyocytes, ultimately resulting in cardiomyocyte cell death. We provide evidence that cardiomyocyte infection is dependent on ACE2 expression and endosomal cysteine protease activity. SARS-CoV-2-infected EHTs displayed typical features of myocarditis, including activation of immune cells, decreased contractile force generation, and cardiomyocyte cell death. Furthermore, autopsy and biopsy samples from four patients with confirmed SARS-CoV-2 infection and clinical myocarditis demonstrated patchy cardiomyocyte infection that was accompanied by myocardial cell death and macrophage accumulation. These findings demonstrate that SARS-CoV-2 can productively infect human cardiomyocytes and establish EHTs as a platform for mechanistic investigation of COVID-19 myocarditis.

## Results

### ACE2 is expressed in human cardiomyocytes

To explore whether human cardiomyocytes might be susceptible to SARS-CoV-2 infection, we examined the expression of angiotensin converting enzyme 2 (ACE2) within the human heart. Previous studies have established that ACE2 serves as a cell-surface receptor for SARS-CoV-2 through interactions with the spike protein in numerous human cell types^15,16^. Immunostaining of human left ventricular myocardial tissue revealed evidence of ACE2 expression in cardiomyocytes (**Fig. 1a**). We observed significant variation in ACE2 expression between individual cardiomyocytes within the same myocardial specimen. ACE2 mRNA was abundantly expressed in the healthy human heart and further increased in the context of chronic heart failure (**Fig. 1b)**. RNA sequencing of human pediatric and adult heart failure specimens revealed robust expression of ACE2 mRNA within the human heart across the spectrum of age (**Fig. 1c**). Consistent with our immunostaining findings, primary human cardiomyocytes obtained from the left ventricle and atria expressed ACE2 mRNA (**Fig. 1d**). These data are consistent with prior single cell and bulk RNA sequencing analyses of human myocardium and suggest that cardiomyocytes might be permissive to SARS-CoV-2 infection^17^.

**Figure 1:**
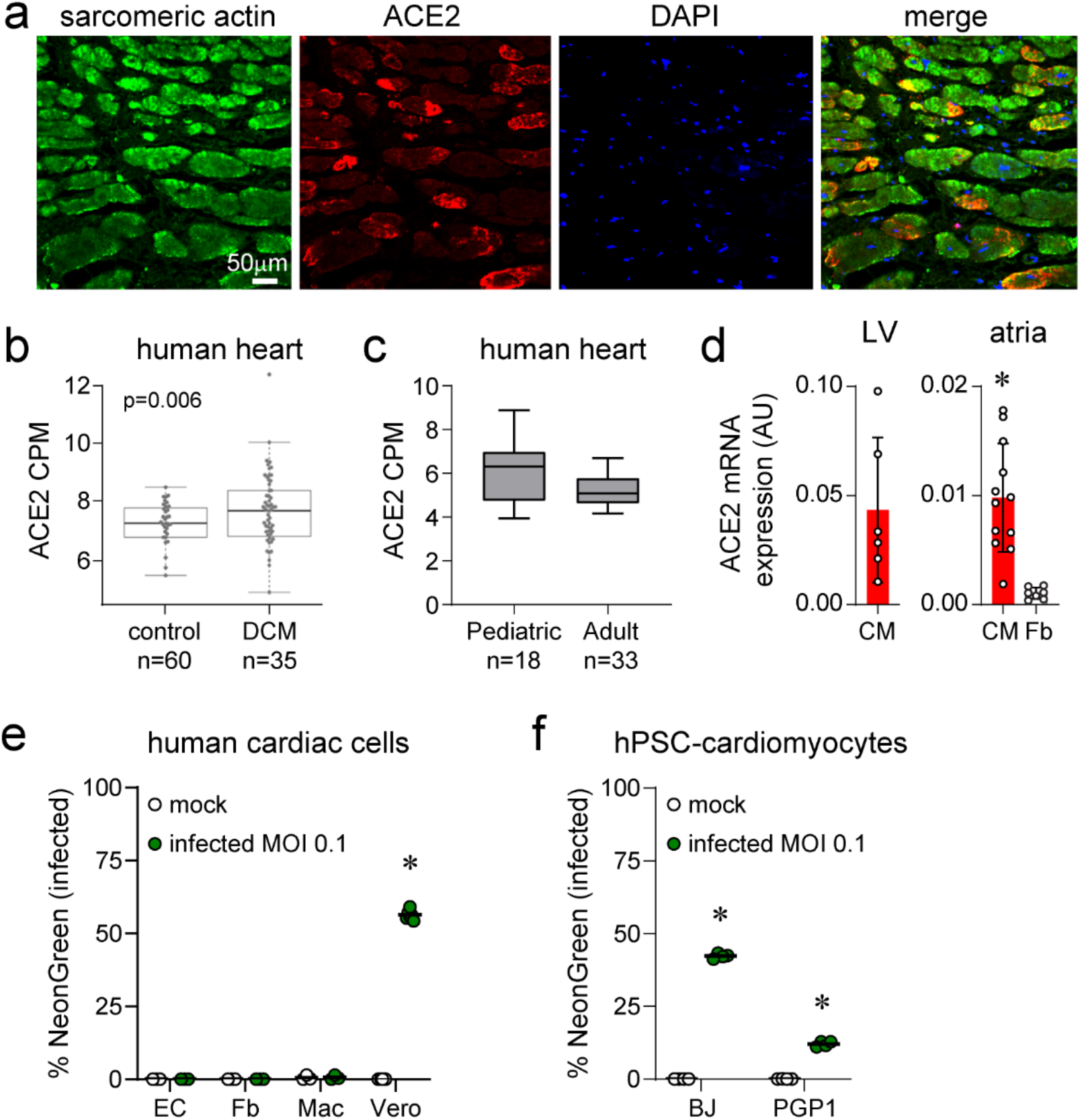
ACE2 is expressed in the human heart and in stem cell derived cardiomyocytes. **a**, Immunohistochemistry of human heart tissue showing ACE2 (red) expression in cardiomyocytes (green, sarcomeric actin). Representative images from 5 analyzed specimens. **b**, RNA sequencing demonstrating ACE2 mRNA expression in myocardial biopsies obtained from adult controls and heart failure patients. Data is displayed as counts per million (CPM). n=18 pediatric, n=33 adult. Each data point indicates an individual sample. n=60 controls, n=35 heart failure. **c**, RNA sequencing demonstrating ACE2 mRNA expression in adult and pediatric heart tissue. Data is displayed as counts per million (CPM). n=18 pediatric, n=33 adult. **d**, Quantitative RT-PCR measurements showing ACE2 mRNA expression in human primary left ventricular (LV), cardiomyocytes (CM), atrial cardiomyocytes, and atrial fibroblasts (Fb). Each data point indicates an individual sample. Error bars denote standard deviation. * p<0.05 compared to atrial fibroblasts (Mann-Whitney test). **e**, Inoculation of primary human endothelial cells (EC), fibroblasts (Fb), and macrophages (Mac) with mock (black) or SARS2-CoV-2-NeonGreen (green, MOI 0.1). Vero cells are included as a positive control. Data is presented as the percent of live cells that express NeonGreen indicating infection. Each data point represents cells isolated from an individual patient sample. Bars denote mean values. * p<0.05 compared to mock infection (Mann-Whitney test). **f**, Inoculation of 2 different hPSC-derived cardiomyocyte lines (BJ, PGP1) with mock (black) or SARS2-CoV-2-NeonGreen (green, MOI 0.1, 1.0). Each data point represents biological replicates. * p<0.05 compared to mock infection (Mann-Whitney test). Bars denote mean values.

To ascertain whether human pluripotent stem cell-derived cardiomyocytes (hPSC-derived CMs) can serve as an appropriate model to study cardiac SARS-CoV-2 infection, we measured ACE2 mRNA expression in hPSC-derived CMs. Quantitative RT-PCR revealed that hPSC-derived CMs abundantly expressed ACE2 mRNA. In contrast, minimal ACE2 mRNA was detected in human dermal fibroblasts, hPSC-derived cardiac fibroblasts, or human fetal cord blood derived-macrophages (**Fig. S1a-c**). Human engineered heart tissues (EHTs) self-assembled between two deformable polydimethylsiloxane (PDMS) posts after mixing cells in an extracellular matrix composed of collagen and matrigel (**Fig. S1d**). EHTs composed of either hPSC-derived CMs and fibroblasts or hPSC-derived CMs, fibroblasts, and macrophages also expressed ACE2 mRNA (**Fig. S1e**). Immunostaining of EHTs confirmed the presence of ACE2 protein specifically in hPSC-derived CMs (**Fig. S1f**). These data suggest that hPSC-derived CMs might be susceptible to SARS-CoV-2 infection and serve as a suitable experimental model to study cardiac manifestations of COVID-19.

### SARS-CoV-2 infects hPSC-derived cardiomyocytes

To determine the susceptibility of different myocardial cell types to SARS-CoV-2 infection, we inoculated various stromal populations with a recombinant SARS-CoV-2 clone containing a NeonGreen fluorescent reporter (SARS-CoV-2-mNeonGreen)^18^. NeonGreen is expressed from a viral subgenomic RNA, indicative of active viral replication. We were unable to detect infection of primary human cardiac fibroblasts, endothelial cells, or macrophages (**Fig. 1e,** gating schemes: **Fig. S2**). hPSC-derived endothelial cells and cardiac fibroblasts were also not susceptible to infection (**Fig. S1g**). In contrast, two independent lines of hPSC-derived cardiomyocytes (hPSC-derived CMs) were permissive to SARS-CoV-2 infection (**Fig. 1f**). Undifferentiated hPSC lines did not demonstrate evidence of infection (**Fig S3**).

To confirm cardiomyocyte tropism, we inoculated various combinations of hPSC-derived CMs, fibroblasts, and macrophages grown in monolayer culture with wild-type SARS-CoV-2 (USA_WA1/2019). We analyzed tissue culture supernatants for production of infectious virus using a Vero cell infection-based focus forming assay, and we measured intracellular viral RNA transcript levels using RT-qPCR at 3 days post-inoculation. These assays revealed the production of infectious virus (**Fig. 2a**) and viral RNA (**Fig. 2b**) selectively in cultures that contained hPSC-derived CMs. Cultures lacking hPSC-derived CMs contained viral loads that were equivalent to media-only controls. A time course of hPSC-derived CM infection showed that cardiomyocytes rapidly produced infectious virus with peak titers observed at day 3 post-inoculation. These kinetics were closely mirrored by SARS-CoV-2-mNeonGreen (**Fig. 2c**).

**Figure 2:**
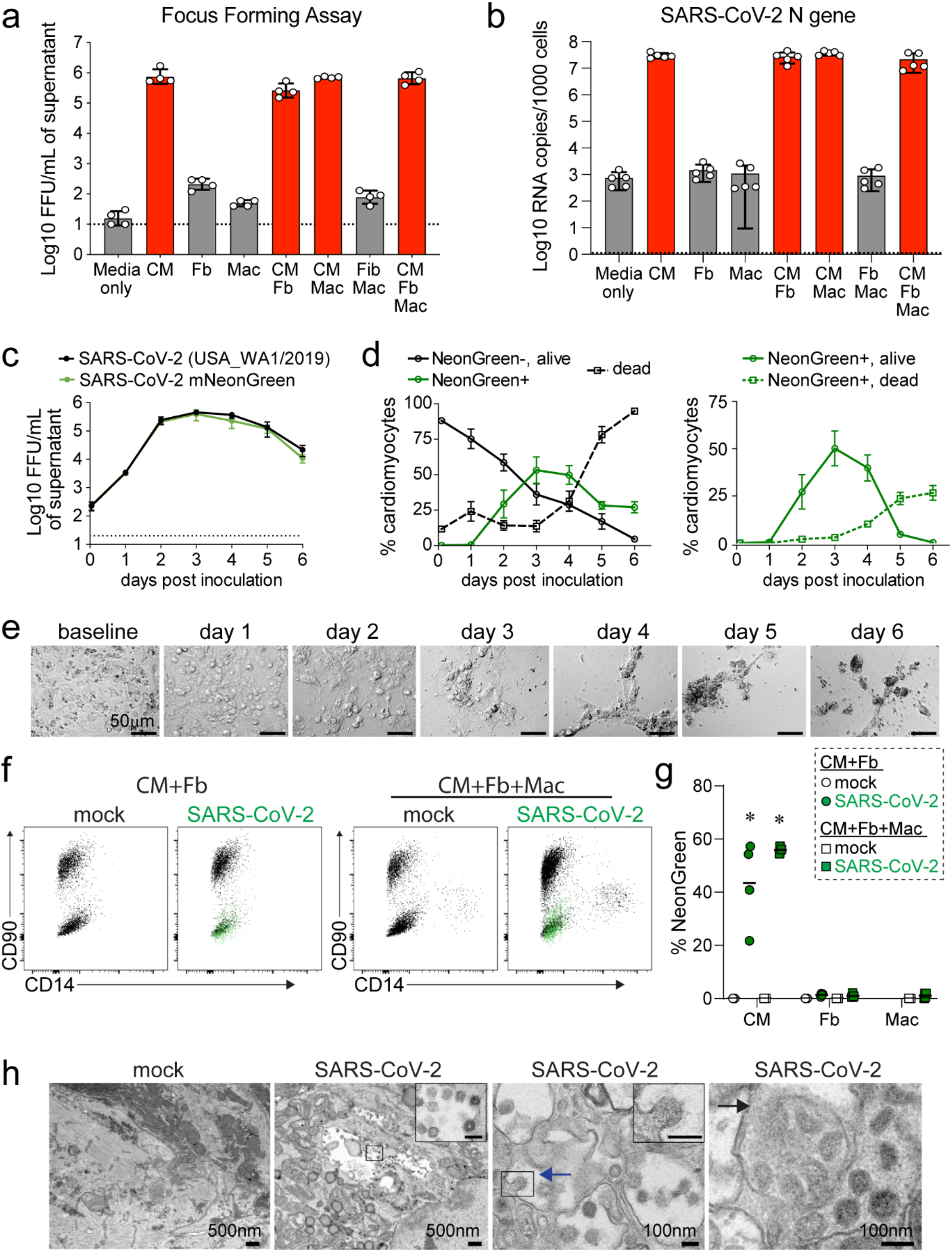
SARS-CoV-2 infects cardiomyocytes. **a**, Focus forming assay demonstrating production of infectious virus from cultures containing hPSC-derived cardiomyocytes (CM), fibroblasts (Fb), and macrophages (Mac) inoculated with SARS-CoV-2 (MOI 0.1). Media only denotes wells that contain no cells. Assays were performed 3 days following inoculation. Dashed line shows the limit of assay detection. **b**, Quantitative RT-PCR showing viral N gene copies in cultures containing CM, Fb, and Macs inoculated with SARS-CoV-2 (MOI 0.1). RNA was collected 3 days after inoculation. Data points indicate individual samples (n=5, a-b). Bars denote the mean value and error bars indicate standard error of the mean. **c**, Focus forming assay measuring infectious SARS-CoV-2 (wild-type, black; mNeonGreen, green) in supernatant of hPSC-derived cardiomyocytes over time following inoculation (MOI 0.1). Dashed line shows the limit of detection. n=4 per experimental group. Error bars denote standard deviation. **d**, Two-dimensional cultures of hPSC-derived cardiomyocytes were inoculated with SARS-CoV-2 (MOI 0.1) and analyzed for viability (Zombie-Violet) and infection (NeonGreen reporter) as a function of time by flow cytometry. Right plot: viability of NeonGreen positive cells. n=4 per experimental group. Error bars denote standard deviation. **e**, Brightfield microscopy showing cytopathic effect (CPE) in hPSC-derived cardiomyocytes infected with SARS-CoV-2 (MOI 0.1). Representative images from 5 individual samples. **f**, Flow cytometry of two-dimensional tissues containing CM and Fb (left) or CM, Fb, and Mac (right) harvested on day 3 following either mock infection or inoculation with SARS-CoV-2 (MOI 0.1). Representative plot from 4 independent samples. Cardiomyocytes (CD90-CD14−) demonstrated prominent NeonGreen fluorescence (green overlay). NeonGreen signal was not detected in fibroblasts (CD90+CD14−) or macrophages (CD90-CD14+). **g**, Quantification of the percent NeonGreen positive cells from 2-dimensional tissues containing hPSC-CMs and fibroblasts or hPSC-CMs, fibroblasts, and macrophages. Data points indicate individual samples (n=4). * p<0.05 compared to mock infection (Mann-Whitney test). **h**, Transmission electron microscopy micrographs of cardiomyocytes in two-dimensional tissues infected with either mock or SARS-CoV-2 (MOI 0.1). Tissues were harvested on day 3 post-inoculation. Virions are readily apparent. Viral budding (blue arrow) and endosomal compartments filled with virions (black arrow) are denoted. Scale bars in insets are 100 nm. Representative image from 4 independent samples.

Using SARS-CoV-2-mNeonGreen, we examined the relationship between viral replication and cell death using flow cytometry. Although the percentage of hPSC-derived CMs that were mNeonGreen-positive peaked at day 3 post-inoculation, significant levels of hPSC-derived CM cell death were not observed until 4-5 days post-inoculation (**Fig. 2d**) indicating that viral infection precedes hPSC-derived CM cell death. SARS-CoV-2-infected cardiomyocytes also displayed characteristics of cytopathic effects. Cellular rounding, clumping, and syncytium formation first were observed on day 3 post-inoculation. Distortion of cellular morphology was evident by day 4 post-inoculation and cultures contained largely dead cells and debris by days 5-6 post-inoculation (**Fig. 2e**).

To verify that cardiomyocytes are the primary target of SARS-CoV-2 in a simulated cardiac environment, we infected two-dimensional tissues assembled with hPSC-derived CMs (80%), fibroblasts (10%), and macrophages (10%) with SARS-CoV-2-mNeonGreen. Flow cytometry performed 3 days following infection revealed mNeonGreen expression only in CD90^−^CD14^−^ TNNT2^+^ cardiomyocytes. mNeonGreen was not detected in CD90^+^ fibroblasts or CD14^+^ macrophages within infected two-dimensional tissues (**Fig. 2f-g, Fig. S4**). These data demonstrate selective viral tropism in cardiomyocytes. Consistent with this conclusion, transmission electron microscopy of infected two-dimensional tissues performed 3 days post-inoculation demonstrated the presence of coronavirus particles within infected hPSC-derived CMs. Micrographs revealed structural features of coronaviruses including the presence of a tri-laminar envelope and characteristic cross-sections through the nucleocapsid^19,20^ (**Fig. 2h**). Virions were identified within perinuclear endosomal-like structures of hPSC-derived CMs. We observed various stages of virion assembly including budding from intracellular membranes. Virions were not detected in mock-infected cardiomyocytes.

### RNA sequencing identified robust viral transcription and the activation of innate immune responses in hPSC-derived cardiomyocytes and two-dimensional tissues

To examine viral transcription and the host immune response to SARS-CoV-2 infection, we performed RNA sequencing. Cultures containing either hPSC-derived CMs, fibroblasts, or macrophages were either mock-infected or inoculated with SARS-CoV-2. We also examined two-dimensional co-culture tissues assembled with 80% cardiomyocytes, 10% fibroblasts, and 10% macrophages. Cells and tissues were harvested on day 3 post-inoculation. Principal component analysis revealed separation between experimental groups consistent with their distinct cellular composition (**Fig. 3a**). Classification of transcript types demonstrated that infected hPSC-derived CMs and two-dimensional tissues comprised of hPSC-derived CMs and fibroblasts or hPSC-derived CMs, fibroblasts, and macrophages contained abundant viral transcripts (**Fig. S5a**). We then assessed the expression of specific viral transcripts by aligning the RNA sequencing data to the SARS-CoV-2 genome and transcriptome. Subgenomic RNAs were identified based on the presence of 5’ leader sequences^21^. We observed robust expression of most SARS-CoV-2 genomic and subgenomic RNAs in infected hPSC-derived CMs and two-dimensional tissues with the exception of ORF7b (**Fig. 3b, Fig. S5b**).

**Figure 3:**
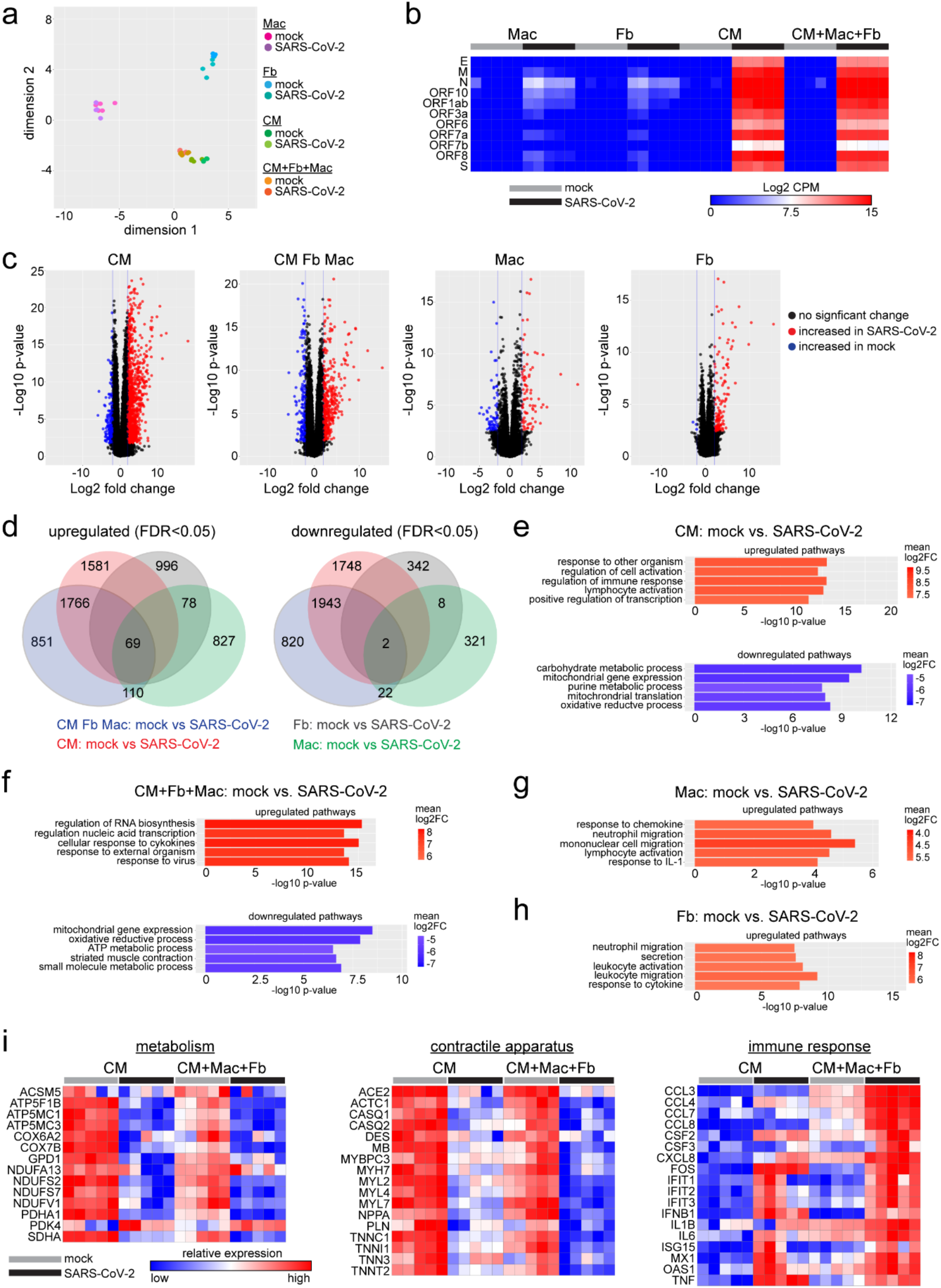
RNA sequencing identified robust viral transcription and activation of innate immune response in hPSC-derived cardiomyocytes and 2D tissues. **a,** MDS plot of RNA sequencing data obtained from mock and SARS-CoV-2 infected (MOI 0.1) hPSC-derived cardiomyocytes (CM), fibroblasts (Fb), macrophages (Mac), and two-dimensional tissues (CM+Fb+Mac) containing all 3 cellular components. Cells and tissues were harvested on day 3 post-inoculation. n=5 per experimental group. **b,** Heatmap of SARS-CoV-2 viral gene expression in each condition. Color scale denotes absolute expression as log2 of counts per million reads (CPM) (blue=0, red=15). **c,** Volcano plots showing differentially expressed genes between mock and SARS-CoV-2 infected conditions. Black: no significant change, Red: upregulated during infection (log2 fold change>2, FDR p-value<0.05), blue: downregulated during infection (log2 fold change<2, FDR p-value<0.05). Data points correspond to individual genes or transcripts. **d,** Venn diagram of genes upregulated and downregulated in each cell type and two-dimensional tissues. Differential expression is based on change relative to corresponding uninfected (mock) samples. **e-h,** GO Pathway analysis of CM (**e**), CM+Fb+Mac (**f**), Mac (**g**) and Fib (**h**) showing top five upregulated (red) and downregulated (blue) pathways in SARS-CoV-2-infected samples compared to mock. Color indicates log2 fold change (log2FC). **i**, Heat maps of selected differentially expressed genes implicated in metabolism (left), contractile apparatus (center) and immune response (right). CM and two-dimensional tissues (CM+Mac+Fb) are displayed. Color scale denotes relative gene expression (high-red, low-blue) across cell types and conditions.

To facilitate differential expression analysis of host genes, we censored viral RNAs from the RNA sequencing computational model. This was necessary given the asymmetric prevalence of viral transcripts across samples. We identified numerous host genes that were differentially regulated upon SARS-CoV-2 infection in each of the examined cell types and two-dimensional tissues (**Fig. 3c**). Conditions that supported viral replication (hPSC-derived CMs and two-dimensional tissues) displayed the greatest overlap in differentially expressed genes. Cell types that did not support viral replication (fibroblasts and macrophages) also demonstrated numerous differentially expressed host genes, indicating that SARS-CoV-2 might elicit changes in host gene expression in the absence of direct viral infection. Notably, host genes differentially expressed in fibroblasts and macrophages exposed to SARS-CoV-2 were largely distinct (**Fig. 3d**). These findings suggest that elements within or on the surface of SARS-CoV-2 virions may serve as pathogen-associated molecular patterns (PAMPs) and stimulate distinct gene expression programs in differing cell types.

GO pathway analysis revealed that infected hPSC-derived CMs and two-dimensional co-culture tissues showed upregulation of genes associated with immune cell activation, stress-induced transcription, and responses to pathogens including viruses. Genes associated with metabolism, oxidative phosphorylation, and mitochondrial function were downregulated by infection. Two-dimensional tissues displayed alterations in other pathways including upregulation of cellular responses to cytokines and downregulation of genes involved in muscle contraction (**Fig. 3e-f**). Host genes differentially expressed in macrophages and fibroblasts were associated with pathways involved in innate immune cell activation, migration, and cytokine responses (**Fig. 3g-h**).

Examination of specific genes differentially regulated in infected hPSC-derived CMs and two-dimensional tissues (**Fig. 3i**) revealed marked reduction in components of the electron transport chain (ATP synthase, mitochondrial cytochrome C oxidase, NADPH dehydrogenase) and key upstream metabolic regulators (glycerol-3-phosphate dehydrogenase, pyruvate dehydrogenase, succinate dehydrogenase complex). *PDK4*, an inhibitor of pyruvate dehydrogenase was upregulated in infected hPSC-derived CMs and two-dimensional tissues. We also observed marked downregulation of numerous components and regulators of the contractile apparatus including cardiac actin, troponin subunits, myosin light and heavy chains, desmin, phospholamban, and calsequestrin in infected two-dimensional tissues. Infected hPSC-derived CMs displayed similar changes, albeit to a lesser extent. *ACE2* expression was diminished in infected cardiomyocytes and two-dimensional tissues.

Infected hPSC-derived CMs and two-dimensional tissues also displayed upregulation of key regulators of innate immunity (**Fig. 3i**). Type I interferon (IFN) activation was apparent by the increased expression of *IFNB1* and numerous IFN stimulated genes including *IFIT1*, *IFIT2*, *IFIT3*, *ISG15*, *MX1*, and *OAS1*. Stress response programs (FOS) and cytokine expression (TNF) were similarly upregulated in these cell types. Consistent with a greater innate immune response in two-dimensional tissues, we found that several chemokines (*CCL3*, *CCL4*, *CCL7*, *CCL8*, and *CXCL8*) and cytokines (*IL1B*, *IL6*, and *CSF3*) were selectively upregulated in infected two-dimensional tissues. Macrophages and fibroblasts contributed to enhanced chemokine and cytokine responses in two-dimensional tissues. *CCL3*, *CCL4*, and *CCL8* were selectively expressed in infected macrophages and *CSF3*, *CXCL8*, *IL1B*, and *IL6* were induced in infected fibroblasts (**Fig. S5c**).

### SARS-CoV-2 entry into hPSC-derived cardiomyocytes is mediated by ACE2 and endosomal cysteine proteases

To elucidate the mechanism(s) by which SARS-CoV-2 enters cardiomyocytes, we examined SARS-CoV-2 infection of hPSC-derived CMs in the presence of well-established entry inhibitors. ACE2 serves as the cell-surface receptor for SARS-CoV-2 in humans^15,16^. Blockade of ACE2 with anti-hACE2 antibody abrogated SARS-CoV-2-NeonGreen infectivity in hPSC-derived CMs as measured by NeonGreen-positivity and viral RNA extracted from the supernatant of infected cultures. The extent of blockade was comparable to treatment with remdesivir, a potent inhibitor of the SARS-CoV-2 RNA-dependent RNA polymerase^22–24^ (**Fig. 4a-b**).

**Figure 4.**
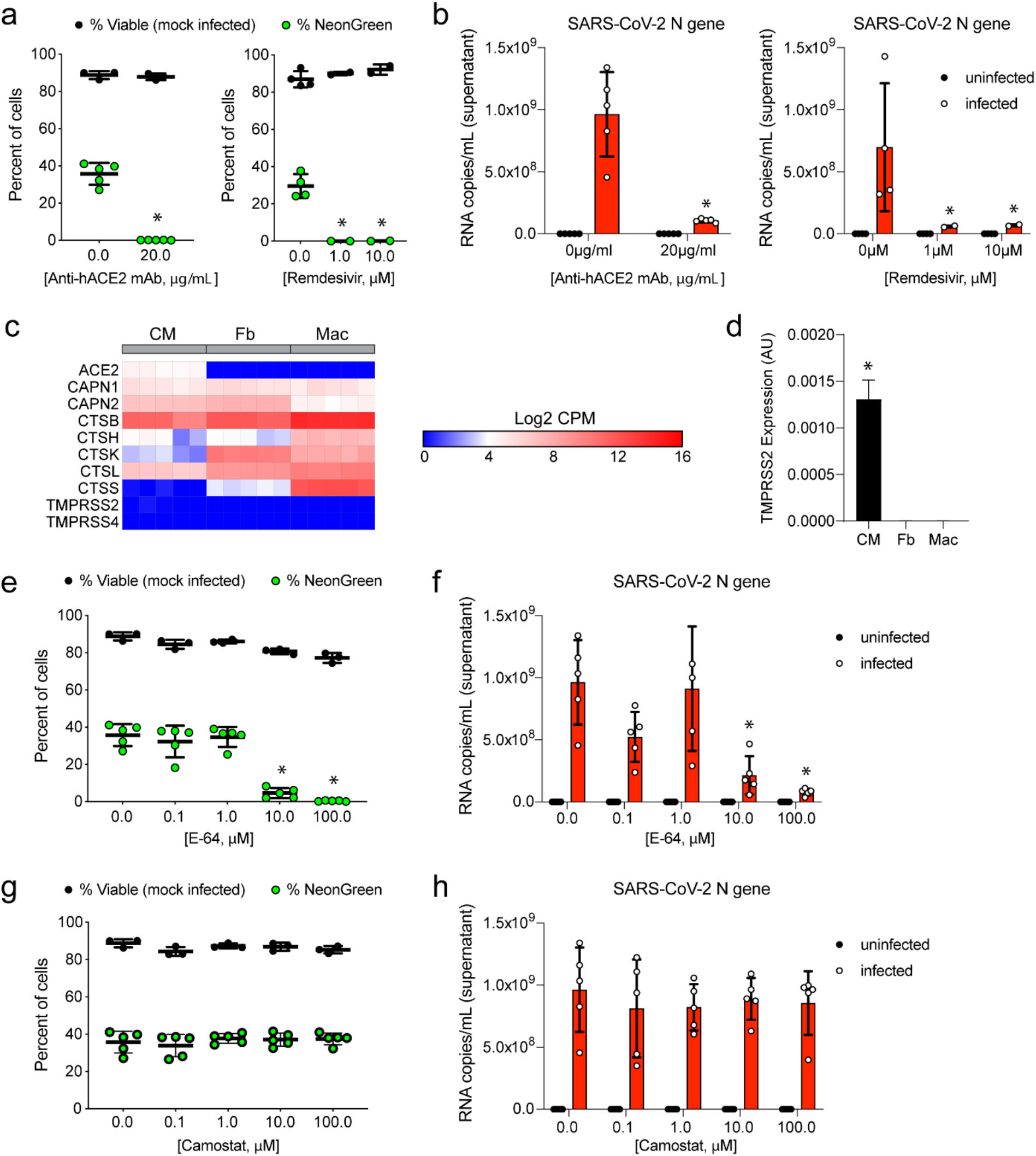
SARS-CoV-2 entry of hPSC-derived cardiomyocytes is mediated by ACE2 and endosomal cysteine proteases. **a-b,** hPSC-derived cardiomyocytes were infected with mock or inoculated with SARS-CoV-2-mNeonGreen (MOI 0.1). Cells were treated with either vehicle control, anti-human ACE2 neutralizing antibody (Anti-hACE2 mAb, left), or remdesivir (inhibitor of RNA-dependent RNA polymerase, right) at the indicated concentrations. Cells were analyzed by flow cytometry on day 3 post-inoculation for viral infection (NeonGreen, green circles) and viability (Zombie-Violet, black circles) (**a**). The presence of viral RNA in the tissue culture supernatant was also quantified by RT-PCR (**b**). Each data point corresponds to an individual sample/experiment, error bars denote standard error of the mean, *p<0.05 compared to infected cells treated with vehicle control (Mann-Whitney test). **c,** Heatmap of host genes implicated in SARS-CoV-2 cell entry in uninfected hPSC-derived cardiomyocytes (CM), fibroblasts (Fb), and macrophages (Mac). Color scale indicates absolute gene expression levels. **d,** Quantitative RT-PCR for *TMPRSS2* in uninfected hPSC-derived cardiomyocytes (CM), fibroblasts (Fb), and macrophages (Mac). n=5 for each cell type, error bars denote standard error of the mean, *p<0.05 compared to other cell populations (Mann-Whitney test). **e-h**, hPSC-derived cardiomyocytes were mock infected or inoculated with SARS-CoV2-mNeonGreen (MOI 0.1). Cells were treated with either vehicle control, endosomal cysteine protease inhibitor E-64 (**e-f**), or serine protease inhibitor camostat (**g-h**) at the indicated concentrations. Cells were analyzed by flow cytometry on day 3 post-inoculation for viral infection (NeonGreen, green circles) and viability (Zombie-Violet, black circles) (**e,g**). Viral RNA in the tissue culture supernatant was quantified by RT-PCR (**f,h**). Each data point corresponds to an individual sample/experiment, error bars denote standard error of the mean, *p<0.05 compared to infected cells treated with vehicle control (Mann-Whitney test).

After binding to ACE2, the SARS-CoV (and SARS-CoV-2) spike protein must undergo proteolytic activation to initiate membrane fusion^25^. Host proteases located at the plasma membrane (TMPRSS2) or within endosomes (cathepsins) most commonly perform this function. The relative contributions of each of these protease families to SARS-CoV-2 infection varies by cell-type^15,25^. RNA sequencing data revealed that hPSC-derived CMs express robust levels of ACE2 and multiple endosomal proteases including cathepsins and calpains (**Fig. 4c**). ACE2 mRNA was not abundantly expressed in either macrophages or fibroblasts. While TMPRSS2 expression was present at the lower limit of detection for RNAseq, we detected low, but measurable levels of TMPRSS2 by RT-qPCR in hPSC-derived CMs, but not in fibroblasts or macrophages (**Fig. 4c-d**).

To determine whether SARS-CoV-2 enters cardiomyocytes through an endosomal or plasma membrane route, we inoculated hPSC-derived CMs with SARS-CoV-2-mNeonGreen and administered either the endosomal cysteine protease inhibitor E-64, which blocks cathepsins, or the serine protease inhibitor camostat mesylate, which blocks TMPRSS2 (and possibly TMPRSS4) ^25^. Notably, E-64 abolished SARS-CoV-2 infection of hPSC-derived CMs as demonstrated by reduced mNeonGreen expression and viral RNA within the supernatant (**Fig. 4e-f**). In contrast, camostat had no effect on cardiomyocyte infection over a range of doses (**Fig. 4g-h**). Thus, SARS-CoV-2 enters cardiomyocytes through an endosomal pathway that requires cathepsin but not TMPRSS2-mediated cleavage.

### EHTs model COVID-19 myocarditis

Myocarditis is characterized by direct viral infection of cardiomyocytes and accumulation of immune cells at sites of active infection or tissue injury^26,27^. To examine whether SARS-CoV-2 infection of cardiomyocytes in a three-dimensional environment mimics aspects of viral myocarditis, we generated EHTs containing either hPSC-derived CMs and fibroblasts or hPSC-derived CMs, fibroblasts, and macrophages. EHTs were seeded in a collagen-Matrigel matrix between two PDMS posts, infected with SARS-CoV-2, and harvested 5 days after inoculation. Hematoxylin and eosin (H&E) staining revealed evidence of tissue injury and increased interstitial cell abundance within the periphery of SARS-CoV-2-infected EHTs (**Fig. 5a**). Immunostaining for the viral nucleocapsid protein demonstrated evidence of prominent infection at the periphery of the EHT. Nucleocapsid staining was localized within hPSC-derived CMs. Staining for CD68 demonstrated macrophage accumulation corresponding to sites of interstitial cell accumulation and viral infection (**Fig. 5b, Fig. S6**). Enrichment of nucleocapsid staining at the periphery of the tissue suggests that viral diffusion might be limited by the three-dimensional EHT environment. Consistent with our immunostaining results, infected EHTs (with and without macrophages) accumulated high levels of viral RNA, as detected by quantitative RT-PCR (**Fig. 5c**). *In situ* hybridization for viral spike sense and antisense RNA was also indicative of viral replication within EHTs (**Fig. 5d, Fig. S7**).

**Figure 5.**
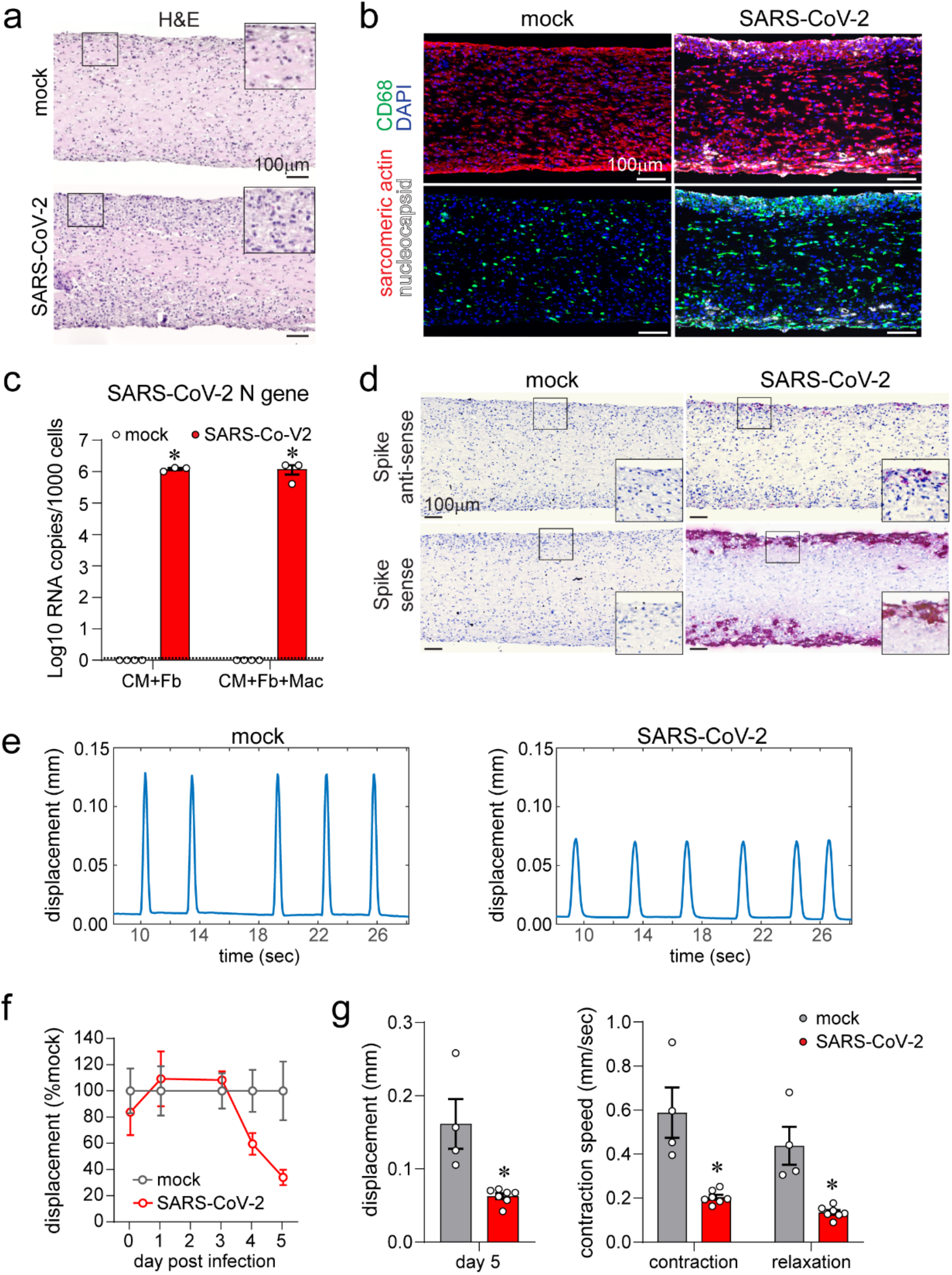
Human engineered heart tissues (EHTs) recapitulate aspects of COVID-19 myocarditis. **a,** Representative hematoxylin and eosin (H&E) stained histology images of three-dimensional EHTs consisting of hPSC-derived cardiomyocytes (CM), fibroblasts (Fb), and macrophages (Macs) 5 days following mock infection or inoculation with SARS-CoV-2 (MOI 0.1). Insets are high magnification images of the boxed areas. Representative images from 4 independent samples. **b**, Immunostaining of mock or SARS-CoV-2 infected three-dimensional EHTs for sarcomeric actin (cardiomyocytes, red), CD68 (macrophages, green), and nucleocapsid protein (white). EHTs were harvested 5 days after inoculation. Blue: DAPI. Images are representative of 4 independent experiments. Representative images from 4 independent samples. **c,** Quantitative RT-PCR of SARS-CoV-2 N gene expression in EHTs consisting of hPSC-derived cardiomyocytes (CM) and fibroblasts (Fb) or hPSC-derived cardiomyocytes, fibroblasts, and macrophages. EHTs were either mock infected or inoculated with SARS-CoV-2 (MOI 0.1) and harvested 5 days after inoculation. Each data point represents individual samples/experiments. Error bars denote standard error of the mean. Bar height represents sample mean. Dotted line: limit of detection. *p<0.05 compared to uninfected control (mock, Mann-Whitney test). **d,** *In situ* hybridization for SARS-CoV-2 ORF1ab RNA sense and anti-sense strands (red) in EHTs 5 days after mock or SARS-CoV-2 infection (MOI 0.1). Hematoxylin: blue. Representative images from 4 independent specimens. Insets are high magnification images of the boxed areas. **e,** Representative spontaneous beating displacement traces for an infected and an uninfected EHT on day 5 post-infection. Videos used to generate these traces can be found in **Supplemental Videos 1 and 2**. **f,** Displacement (relative to uninfected mock condition) generated by spontaneous beating of EHTs as a function of time following inoculation with SARS-CoV-2 (MOI 0.1). Each data point represents a mean value from 4-7 independent samples (2 independent experiments), error bars denote standard error of the mean. **g,** Quantification of absolute displacement (left) and contraction speed (right) generated by spontaneous beating of EHTs 5 days following mock or SARS-CoV-2 infection (MOI 0.1). Each data point denotes an individual EHT, bar height corresponds to mean displacement, error bars represent standard error of the mean, *p<0.05 compared to mock (Mann-Whitney test).

### SARS-CoV-2 infection causes contractile dysfunction and cell death

Reduced left ventricular systolic function has been reported in severe cases of COVID-19 myocarditis^28^. Therefore, we examined the effect of SARS-CoV-2 infection on cardiomyocyte contractile function in EHTs. EHTs were seeded between two deformable PDMS posts of known stiffness. As the tissue contracts, it displaces the posts, and by tracking the displacement of the posts as a function of time, we calculated the speed of contraction and relaxation. The average peak displacement was calculated for each spontaneously contracting tissue over the course of at least 60 seconds (**Supplemental Movies 1-2**).

EHTs consisting of hPSC-derived CMs and fibroblasts were assembled and allowed to mature for 7 days prior to infection. EHTs were inoculated with SARS-CoV-2, and contractile function was analyzed daily. From days 0 to 3 post infection, the average maximal displacement generated during beating did not differ between the mock and SARS-CoV-2-infected tissues. However, on days 4 to 5 post infection, the SARS-CoV-2 inoculated tissues showed reduced contraction relative to the mock-infected tissues (**Fig. 5e-f**). On day 5 after inoculation, the maximal displacement produced during contraction by the SARS-CoV-2 inoculated tissues was markedly lower than mock infected-tissues. Moreover, the tissues show reduced speed of contraction and relaxation, consistent with systolic dysfunction (**Fig. 5g**).

### Mechanisms of reduced EHT contractility

To examine whether cardiomyocyte cell death might serve as a mechanism explaining reduced EHT contractility on days 4 to 5 post inoculation, we performed TUNEL staining. Consistent with the temporal course of SARS-CoV-2 cardiomyocyte infection and cell death in our two-dimensional hPSC-CM cultures (**Fig. 2d**), we observed increased numbers of TUNEL positive cardiomyocytes in SARS-CoV-2 infected EHTs on day 5 post infection (**Fig. 6a-b**). Our RNA sequencing data suggest that other mechanisms also may contribute to reduced EHT contractility, including decreased expression of genes important for sarcomere function and metabolism as well as activation of host immune responses (**Fig. 3i**). Consistent with the possibility that disrupted sarcomere gene expression might contribute to reduced EHT contractility, immunostaining of hPSC-derived CMs infected with SARS-CoV-2 revealed evidence of sarcomere loss 3 days following infection (**Fig. 6c**), a time point that preceded cell death. Furthermore, immunostaining of EHTs demonstrated loss of Troponin T expression in infected cardiomyocytes. (**Fig. 6d-e**). Thus, the reduction in contractile function may be multifactorial with contributions from virus-induced cardiomyocyte cell death and loss of sarcomere elements.

**Figure 6.**
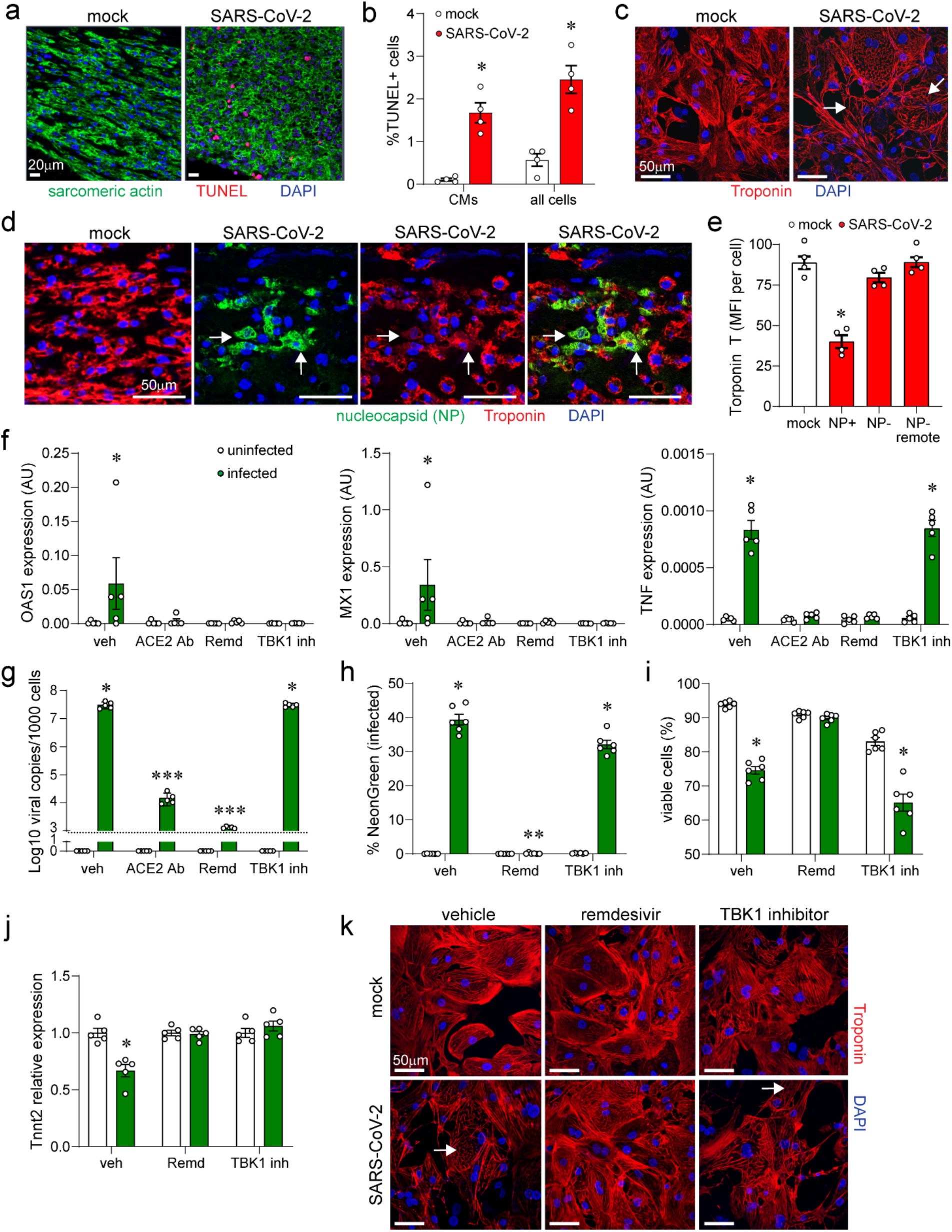
Mechanisms of reduced EHT contractility. **a**, Combined immunostaining for cardiomyocytes (cardiac actin, green) and TUNEL staining (red) of EHTs (CM+Fb+Mac) 5 days after mock or SARS-CoV-2 infection (MOI 0.1). DAPI: blue. Representative images from 4 independent experiments. **b,** Quantification of cell death (percent of TUNEL-positive cells) in areas of viral infection. Each data point denotes an individual EHT, bar height corresponds to the mean, error bars represent standard error of the mean, *p<0.05 compared to mock (Mann-Whitney test). **c**, Immunostaining of hPSC-derived cardiomyocytes for Troponin T (red) 3 days after inoculation with mock control or SARS-CoV-2-NeonGreen (MOI 0.1). Blue: DAPI. Arrows denote areas of sarcomere disassembly. **d**, Immunostaining of EHTs for Troponin T (red) and SARS-CoV-2 nucleocapsid (green) 5 days after inoculation with mock control or SARS-CoV-2- NeonGreen (MOI 0.1). Blue: DAPI. Arrows denote SARS-CoV-2 nucleocapsid positive cells with reduced Troponin T staining. **e**, Quantification of Troponin T staining in mock (white) and SARS-CoV-2 (red) infected EHTs. NP: nucleocapsid. Data is presented as mean florescence intensity (MFI). MFI was measured in infected (NP+) cardiomyocytes and uninfected (NP−) cardiomyocytes located proximal or remote to areas of infection. Each data point denotes an individual EHT, bar height corresponds to the mean, error bars represent standard error of the mean, *p<0.05 compared to mock (Mann-Whitney test). **f**, Quantitative RT-PCR measuring OAS1, MX1, and TNF mRNA expression in hPSC-derived cardiomyocytes 3 days after inoculation with mock control (white) or SARS-CoV-2 (green, MOI 0.1). Cells were treated with vehicle, ACE2 antibody (ACE2 Ab) (20μg/ml), remdesivir (10μM), or TBK inhibitor (MRT67307, 10μM). Each data point denotes a biologically unique sample, bar height corresponds to the mean, and error bars indicate standard error of the mean. * p<0.05 compared to mock control. **g**, Quantitative RT-PCR of SARS-CoV-2 N gene expression in hPSC-derived cardiomyocytes that were either mock infected (white) or inoculated with SARS-CoV-2 (green, MOI 0.1) and harvested 3 days after inoculation. Cells were treated with vehicle, ACE2 Ab (20μg/ml), remdesivir (10μM), or TBK1 inhibitor (MRT67307, 10μM). Each data point represents individual samples. Error bars denote standard error of the mean. Bar height represents sample mean. Dotted line: limit of detection. *p<0.05 compared to uninfected control. ***p<0.05 compared to uninfected control and vehicle infected (mock, Mann-Whitney test). **h-i**, Flow cytometry measuring the percent of infected (h) and viable (i) hPSC-derived cardiomyocytes following either mock infection (white) or inoculation with SARS-CoV-2 (green, MOI 0.1). Cells were harvested and analyzed 3 days after inoculation. Cells were treated with vehicle, remdesivir (10μM), or TBK1 inhibitor (MRT67307, 10μM). Each data point represents individual samples. Error bars denote standard error of the mean. Bar height represents sample mean. *p<0.05 compared to uninfected control. **p<0.05 compared to vehicle infected (mock, Mann-Whitney test). **j,** Quantitative RT-PCR measuring TNNT2 mRNA expression in hPSC-derived cardiomyocytes 3 days after inoculation with mock control (white) or SARS-CoV-2 (green, MOI 0.1). Cells were treated with vehicle, ACE2 Ab (20μg/ml), remdesivir (10μM), or TBK inhibitor (MRT67307, 10μM). Each data point denotes a biologically unique sample, bar height corresponds to the mean, and error bars indicate standard error of the mean. * p<0.05 compared to mock control. **k**, Immunostaining of hPSC-derived cardiomyocytes for Troponin T (red) 3 days after inoculation with mock control or SARS-CoV-2-NeonGreen (MOI 0.1). hPSC-derived cardiomycoytes were treated with vehicle, remdesivir (10μM) or TBK inhibitor (MRT67307, 10μM). Blue: DAPI. Arrows denote areas of sarcomere disassembly. Merged images can be found in Fig. S8.

We then examined the mechanistic relationship between cardiomyocyte infection, inflammatory signaling, sarcomere breakdown, and cell death. Inhibition of viral entry (ACE2 neutralizing antibody) or viral replication (remdesivir) was sufficient to prevent type I IFN and TNF expression following SARS-CoV-2 infection (**Fig. 6f-g**). Remdesivir similarly reduced inflammatory gene expression in 3D EHTs (**Fig. S8a-b**). These data establish that viral infection represents the upstream driver of inflammation in our model system.

To examine the impact of cardiomyocyte inflammatory signaling on cardiomyocyte cell death, sarcomere gene expression, and sarcomere structure, we focused on inhibiting viral nucleic acid sensing. TBK1 (TANK-binding kinase 1) is an essential mediator of numerous nucleic acid sensing pathways including RIG-I, MAVS, STING, and TLRs ^29,30^. Inhibition of TBK1 activity was sufficient to reduce type I IFN activity (primary inflammatory signature identified in infected cardiomyocytes, **Fig 3i**) without impacting viral load or cardiomyocyte infectivity (**Fig. 6f-h**). Inhibition of TBK1 activity during SARS-CoV-2 cardiomyocyte infection had no impact on cardiomyocyte cell death (**Fig. 6i**). While TBK1 inhibition prevented reductions in TNNT2 and MYH7 mRNA expression following cardiomyocyte SARS-CoV-2 infection, sarcomere breakdown remained prevalent in infected cardiomyocytes treated with the TBK1 inhibitor. In contrast, remdesivir prevented both reductions in TNNT2 and MYH7 mRNA expression and sarcomere loss following SARS-CoV-2 infection (**Fig. 6j-k, Fig. S8c-d)**. These data indicate that SARS-CoV-2 elicits an inflammatory response in cardiomyocytes that is at least partially dependent on viral nucleic acid sensing and TBK1 signaling. However, TBK1-dependent cardiomyocyte inflammation does not appear responsible for sarcomeric disassembly or cardiomyocyte cell death. These findings do not rule out the possibility that other inflammatory pathways or cross-talk between infected cardiomyocytes and immune cells contributes to reduced EHT contractility.

### Evidence of cardiomyocyte infection in COVID-19 myocarditis

To validate the myocarditis phenotype generated by SARS-CoV-2 infection of the EHT model, we obtained autopsy and endomyocardial biopsy specimens from four subjects with confirmed SARS-CoV-2 infection and clinical diagnoses of myocarditis. Evidence of myocardial injury (elevated troponin) and left ventricular systolic dysfunction were present in each case (**Table 1**). Coronary angiography demonstrated no evidence of luminal stenosis or thrombosis. The presence of SARS-CoV-2 RNA from nasopharyngeal samples was confirmed by clinical diagnostic testing or post-mortem.

Postmortem microscopic examination of the left ventricular myocardium demonstrated areas of cardiomyocyte necrosis and degenerative vacuolization of cardiomyocyte cytoplasm were noted accompanied by a mixed mononuclear cell infiltrate (**Fig. 7a**). These changes are distinct from postmortem autolytic changes. Examination of the coronary arteries from the COVID-19 myocarditis autopsy cases demonstrated non-obstructive mild atherosclerotic changes, consistent with the angiogram findings. There was no evidence of microvascular injury or thromboembolic events. Two autopsy heart samples from subjects with metastatic carcinoma and an inherited neurodegenerative disease with similar tissue procurement times were included as negative controls.

**Figure 7.**
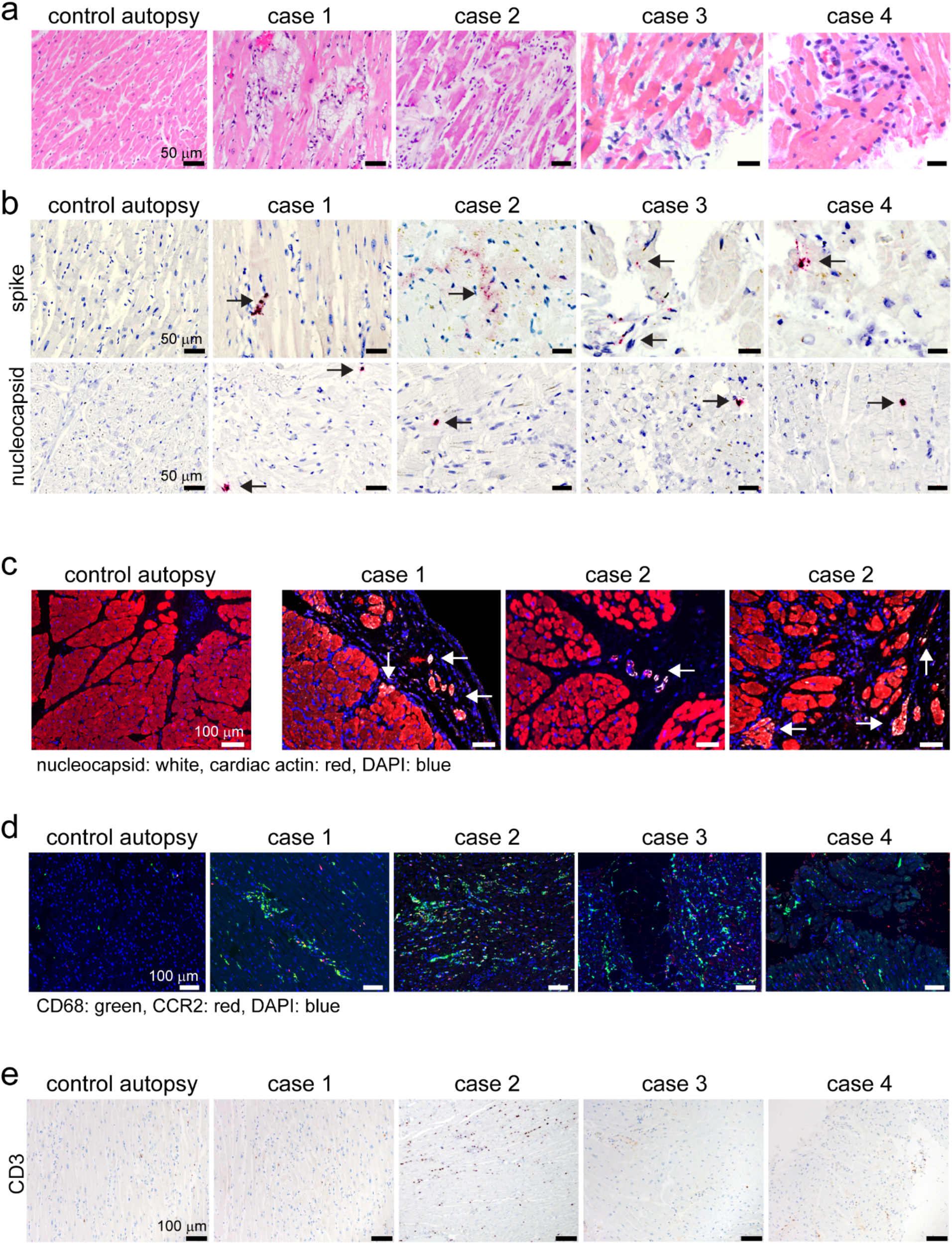
Human autopsy and endomyocardial tissue from patients with suspected COVID-19 myocarditis show evidence of SARS-CoV-2 cardiomyocyte infection. **a**, Hematoxylin and eosin staining of cardiac autopsy (anterior left ventricular wall) and biopsy samples (right ventricular septum) from subjects without COVID-19 (control case) and patients with a clinical diagnosis of COVID-19 myocarditis (case 1-4). **b,** *In situ* hybridization of cardiac autopsy and biopsy tissue for SARS-CoV-2 spike and nucleocapsid RNA (red) showing evidence of viral infection. Hematoxylin: blue. Arrows denotes viral RNA staining in cells with cardiomyocyte morphology. **c**, Immunostaining of control and COVID-19 myocarditis cardiac autopsy tissue for SARS-CoV-2 nucleocapsid (white) and cardiac actin (red). DAPI: blue. Arrows denotes nucleocapsid staining in cardiomyocytes. **d,** Immunostaining of control and COVID-19 myocarditis cardiac autopsy and biopsy tissue for CD68 (green) and CCR2 (red). DAPI: blue. **e,** Immunostaining of control and COVID-19 myocarditis cardiac autopsy and biopsy tissue for CD3 (brown). Hematoxylin: blue.

RNA *in situ* hybridization for SARS-CoV-2 spike and nucleocapsid genes revealed evidence of viral RNA within the myocardium of each COVID-19 myocarditis subject. Viral transcripts were located in cytoplasmic and perinuclear locations within cells that were morphologically consistent with cardiomyocytes (**Fig. 7b, Fig. S9a-b**). Viral transcripts also were identified in airway epithelial cells within the lung of this subject and other myocardial cell types including perivascular adipocytes and pericytes (**Fig. S9c**). Immunostaining for the nucleocapsid protein further demonstrated presence of viral protein in cardiomyocytes **Fig. 7c**). The COVID-19 myocarditis immune cell infiltrate was characterized by accumulation of a mixed population of CCR2^−^ and CCR2^+^ macrophages within injured areas of the myocardium (**Fig. 7d**). Minimal evidence of T-cell infiltration was noted (**Fig. 7e**). Macrophage abundance was highest in areas that demonstrated evidence of cardiomyocyte injury as depicted by complement deposition (C4d staining), a pathological marker of cardiomyocyte cell death^31–33^ (**Fig. S9d**). Together, these observations provide initial pathological evidence that SARS-CoV-2 infects the human heart and may contribute to cardiomyocyte cell death and myocardial inflammation that is distinct from lymphocytic myocarditis.

## Discussion

Cardiac manifestations of COVID-19 including myocardial injury (elevated troponin), reduced left ventricular systolic function, and arrhythmias are increasingly recognized as important determinants of morbidity and mortality^3,8^. Moreover, imaging studies have uncovered evidence of chronic myocardial pathology among both hospitalized and ambulatory patients, suggesting the potential for long-term consequences of COVID-19^5,6^. Little is understood regarding the etiology of these conditions. To gain insight into the cardiac complications of COVID-19, we developed a human EHT system that recapitulates many features of SARS-CoV-2 induced myocarditis. We provide evidence that SARS-CoV-2 infects hPSC-derived CMs, resulting in reduced metabolic and contractile apparatus gene expression, sarcomeric disassembly, inflammatory signaling, and cell death. Viral entry was ACE2-dependent and relied on endosomal cysteine protease activity. Our findings are consistent with a recent report suggesting that SARS-CoV-2 infects human cardiac slices and hPSC-derived CMs in an ACE2 and cathespin dependent manner^34^. We extend these observations to show that cardiomyocytes supported viral replication, rapidly produced infectious virions, activated type I IFN signaling, and displayed cytopathic features seen with coronavirus infection. Infected EHTs demonstrated reduced contractile force, sarcomere disassembly, and pathological evidence of myocarditis including macrophage activation. Examination of human autopsy and endomyocardial biopsy tissue from individuals diagnosed with COVID-19 myocarditis revealed similar findings of cardiomyocyte infection, cell death, and macrophage infiltration.

Whether cardiac manifestations of COVID-19 are a result of viral infection or exuberant systemic inflammation remains a debated topic. It is also possible that microvascular thrombosis may contribute to cardiac events. We provide evidence that SARS-CoV-2 readily infects and replicates within human cardiomyocytes, indicating that viral infection likely contributes to the pathogenesis of COVID-19 myocarditis. SARS-CoV-2 was unable to replicate in other cell types found within the left ventricular myocardium including cardiac fibroblasts, endothelial cells, and macrophages. It remains possible that SARS-CoV-2 could also infect other cardiac cell types that are difficult to isolate from the human heart or produce through directed differentiation protocols, such as pericytes and endocardial cells. Despite these limitations, our findings clearly demonstrate that cardiomyocytes are a target of SARS-CoV-2 infection.

Consistent with this conclusion, examination of human autopsy and endomyocardial biopsy tissue obtained from four patients with clinical diagnoses of COVID-19 myocarditis revealed evidence of myocardial SARS-CoV-2 RNA and protein predominately within cardiomyocytes and accumulation of macrophages in areas surrounding myocardial injury. These findings are consistent with prior reports highlighting infiltration of monocytes, lymphocytes, and plasma cells in an endomyocardial biopsy specimen from a patient with suspected COVID-19 myocarditis^35^ and viral RNA within the myocardium of COVID-19 autopsy specimens^36^. It is important to note that the human specimens examined in this study differ substantially from published autopsy series that did not include subjects with cardiac manifestations ^3,37^. Here, we exclusively focused on subjects with active COVID-19 infection and clinical evidence of myocarditis based on echocardiography and clinical presentation.

Numerous studies have reported that extrapulmonary cell types are susceptible to SARS-CoV-2 infection^38–40^. This broader cellular tropism appears to be dictated by ACE2 expression and the ability of the virus to gain access to extrapulmonary tissues. Among cell types within the heart, cardiomyocytes and pericytes express ACE2 mRNA^17^. Our immunostaining studies of human heart samples and EHTs suggest that ACE2 protein is most prominently expressed in cardiomyocytes. Using a human ACE2 neutralizing antibody, we demonstrated that ACE2 is essential for SARS-CoV-2 to infect cardiomyocytes. It is not yet clear at what stage during the course of their differentiation developing cardiomyocytes become permissive to SARS-CoV-2 infection. Furthermore, it remains to be explored whether cardiomyocyte maturation, remodeling, and/or subset diversification might impact vulnerability or host responses to viral infection. This possibility is supported by the heterogeneous expression of ACE2 in the human heart and hPSC-derived CMs, and may explain why pre-existing cardiovascular disease represents such a strong risk factor for COVID19 mortality. Consistent with this idea, ACE2 expression is increased in heart failure^28,41^. hPSC-derived CMs and EHTs offer an opportunity to address these important questions. Indeed, a previous study demonstrated that hPSC-derived cells (including cardiomyocytes) can support SARS-CoV-2 infection^40^. Whether SARS-CoV-2 enters the heart through hematological seeding and/or direct extension from the pleural cavity remains unknown.

Emerging data suggests that SARS-CoV-2 can enter cells either directly through the plasma membrane using TMPRSS2 or through endosomal pathways via cathepsins, and that the utilization of these pathways is cell-type dependent^15,16^. We demonstrate that SARS-CoV-2 depends almost exclusively upon the endosomal pathway for entry into hPSC-derived CMs, which is consistent with the greater abundance of transcripts for endosomal cathepsins relative to TMPRSS2 and TMPRSS4 transcripts. As such, hPSC-derived CMs may be informative for the development and testing of drugs that target viral cell entry. Based on this, we predict that compounds targeting endosomal viral entry would have greater efficacy in preventing COVID-19 cardiac infection than those targeting TMPRSS2 or related proteases.

EHTs provided an opportunity to gain mechanistic insights into the relationship between cardiomyocyte infection, myocardial inflammation, and contractile dysfunction. Infection of EHTs resulted in generation of inflammatory mediators, reduced ACE2 expression, cardiomyocyte cell death, disrupted sarcomeric structure, and changes in sarcomeric and metabolic gene expression. Each of these mechanisms likely contribute to reduced EHT contractility. We demonstrated that cardiomyocyte infection is essential for inflammatory gene expression, sarcomere loss, and cell death. Intriguingly, blockade of viral nucleic acid sensing pathways (TBK1 inhibitor) blunted reductions in sarcomere gene expression. However, sarcomere disassembly in infected cells and cardiomyocyte cell death was not impacted. These findings highlight the central role of cardiomyocyte infection and suggest that targeting pathways that are responsible for sarcomere breakdown may improve outcomes in patients with cardiac complications of COVID-19. However, our findings do not exclude an important role for inflammation in COVID-19 myocarditis. The relevance of ACE2 downregulation in infected cardiomyocytes will require further clarification as *Ace2*^−/−^ mice display left ventricular systolic dysfunction and heart failure^42^.

Effectively targeting the inflammatory response to SARS-CoV-2 infection in the heart will require a careful dissection of the cell types involved and identification of key effector pathways. Macrophages and fibroblasts appear to contribute to the inflammatory response. Despite low levels of ACE2 expression and resistance to SARS-CoV-2 infection, macrophages and fibroblasts each generated inflammatory chemokines and cytokines in response to exposure to SARS-CoV-2 in 2D culture, a response likely driven by recognition of viral RNAs and/or proteins. The addition of fibroblasts and macrophages to 2D engineered heart tissues exaggerated production of immune-related mRNAs following exposure to SARS-CoV-2. While this response could be a result of direct recognition of viral RNAs and proteins by fibroblasts and macrophages, additional mechanisms of macrophage and fibroblast activation should be considered such as communication between infected cardiomyocytes and adjacent macrophages and fibroblasts through either the production of soluble mediators or intercellular transfer via gap junctions. Future studies are necessary to dissect the cellular mechanisms and signaling pathways that initiate and potentiate local inflammation between infected cardiomyocytes and surrounding macrophages, fibroblasts, and other cell types.

EHTs have been considered as models for dilated and hypertrophic cardiomyopathies given their genetic etiologies^43,44^. Here, we developed a human EHT model of viral myocarditis. EHTs infected with SARS-CoV-2 recapitulated several features of viral myocarditis including cardiomyocyte infection, cell death, inflammation, and contractile dysfunction. These EHTs contained hPSC-derived CMs, monocyte-derived macrophages, and fibroblasts to mirror some of the cellular and extracellular constituents of the human myocardium. To our knowledge, macrophages have not been previously incorporated into human EHTs. Given recent advances in stem cell differentiation protocols, incorporation of tissue resident macrophages, endothelial cells, and cardiac fibroblasts is on the horizon and could further strengthen this model system^45,46^. Moreover, it is possible to genetically modify each of these cellular components to gain mechanistic insights into the pathogenesis of various diseases including COVID-19 myocarditis. These observations add to an emerging field that have successfully utilized EHT systems for testing of compounds for cardiotoxicity and efficacy^47,48^. From a broader perspective, our findings are consistent with the emerging utility of human organoid systems in the investigation of SARS-CoV-2^38,39,49,50^.

Our study is not without limitations, including limited autopsy and biopsy tissue availability, inherent immaturity of hPSC-derived CMs, and incomplete representation of human myocardial cell types included in EHTs. Nonetheless, our experiments support the conclusion that SARS-CoV-2 infection of cardiomyocytes and resultant myocardial injury and inflammation likely promote or contribute to cardiac manifestations. We provide evidence that human EHTs recapitulate many features of COVID-19 myocarditis, demonstrate that SARS-CoV-2 infection of EHTs can produce multiscale changes spanning from the molecular to functional levels, and show that EHTs serve as useful tools for dissecting mechanisms that contribute to cardiac pathology.

## Methods

### Biosafety

All aspects of this study were approved by the office of Environmental Health and Safety at Washington University School of Medicine prior to the initiation of this study. Work with SARS-CoV-2 was performed in a BSL-3 laboratory by personnel equipped with powered air purifying respirators (PAPR).

### Viruses

The 2019n-CoV/USA_WA1/2019 isolate of SARS-CoV-2 was obtained from the United States Centers for Disease Control (CDC). The mNeonGreen SARS-CoV-2 virus was published recently ^18^. Infectious stocks were grown by inoculating Vero CCL81 cells and collecting supernatant upon observation of cytopathic effect. Debris was removed by centrifugation and passage through a 0.22 μm filter. Supernatant was then aliquoted and stored at −80°C. All infections were performed at a multiplicity of infection (MOI) of 0.1. The mNeonGreen SARS-CoV-2 virus stock used in this study was subjected to deep sequencing using ARTIC^51^ and found to have a mutation in the furin cleavage site (positions 23606-23608 of NC_045512.2) at a combined frequency of 31%.

### Focus forming assay

Vero E6 cells were seeded at a density of 4×10^4^ cells per well in flat-bottom 96-well tissue culture plates. The following day, media was removed and replaced with 100 μL of 10-fold serial dilutions of the material to be titered. Two hours later, 135 μL of methylcellulose overlay was added. Plates were incubated for 48 h, then fixed with 4% paraformaldehyde (final concentration) in phosphate-buffered saline for 20 min, followed by permeabilization with saponin-containing buffer. Plates were incubated overnight at 4°C in 100 μL of permeabilization buffer containing 1 μg/mL of the CR3022 anti-spike monoclonal antibody^52^. Following washing, 50 μL of goat anti-human secondary antibody conjugated to HRP (Sigma AP504P), diluted 1:1000 in permeabilization buffer, was added for 2 hours at room temperature with shaking. Foci were stained with 50 μL of KPL Trueblue (SeraCare), then scanned and automatically quantitated on a Biospot plate reader (Cellular Technology Limited).

### Quantitative RT-PCR

RNA was extracted using the MagMax mirVana Total RNA isolation kit (Thermo Scientific) on the Kingfisher Flex extraction robot (Thermo Scientific). RNA was reverse transcribed and amplified using the TaqMan RNA-to-CT 1-Step Kit (ThermoFisher). Reverse transcription was carried out at 48°C for 15 min followed by 2 min at 95°C. Amplification was accomplished over 50 cycles as follows: 95°C for 15 s and 60°C for 1 min. Copies of SARS-CoV-2 N gene RNA in samples were determined using a previously published assay^53^. Briefly, a TaqMan assay was designed to target a highly conserved region of the N gene (Forward primer: ATGCTGCAATCGTGCTACAA; Reverse primer: GACTGCCGCCTCTGCTC; Probe: /56-FAM/TCAAGGAAC/ZEN/AACATTGCCAA/3IABkFQ/). This region was included in an RNA standard to allow for copy number determination down to 10 copies per reaction. The reaction mixture contained final concentrations of primers and probe of 500 and 100 nM, respectively. For host genes (ACE2, TMPRSS2, OAS1, MX1, TNF), RNA was reverse-transcribed using the High Capacity cDNA Reverse Transcription Kit (ThermoFisher) and amplified using SYBR Green system (ThermoFisher) with beta2 microglobulin as an internal reference gene.

### Quantification of ACE2 expression in RNA-Seq

We performed a differential gene expression analysis using R package DESeq2^54^ on the Care4DCM cohort^55^. The DESeq2 package utilizes negative binominal distribution to model the distribution of RNA-Seq reads, and uses Wald test to calculate p-values^54^. The Care4DCM cohort included 60 DCM patients and 35 controls^55^. Of the 60 DCM patients, 52 received ACEI and 8 received ARB. The myocardial biopsies were extracted from the LV apex by heart catheterization and preserved in liquid nitrogen following standardized protocols. The RNA was extracted from the cardiac tissues using Allprep Kits (Qiagen, Düsseldorf, Germany), then the RNA sequencing was carried out using the TrueSeq RNA Sample Prep Kit (Illumina, San Diego, California, USA)^55^.

### Human atrial tissue samples

The study protocol involving human tissue samples was approved by the ethics committees of the Medical Faculty of Heidelberg University (Germany; S-017/2013). Written informed consent was obtained from all patients and the study was conducted in accordance with the Declaration of Helsinki. Tissue samples of right atrial appendages (RAA) were obtained from patients undergoing open heart surgery for coronary artery bypass grafting or valve repair /replacement in the local heart surgery department.

### Atrial and ventricular cardiomyocyte isolation

After excision, tissue samples were immediately placed into preoxygenated transport solution (100 mM NaCl, 10 mM KCl, 1.2 mM KH2PO4, 5 mM MgSO4, 50 mM taurine, 5 mM 3-[N-morpholino] propane sulfonic acid [MOPS], 30 mM 2,3-butanedione monoxime and 20 mM glucose, pH 7.0 with NaOH) and subjected to cardiomyocyte isolation within 15 minutes. RAA tissue samples were dissected into small pieces and rinsed 3 times in Ca2+-free Tyrode’s solution (100 mM NaCl, 10 mM KCl, 1.2 mM KH2PO4, 5 mM MgSO4, 50 mM taurine, 5 mM 3-(N-morpholino) propane sulfonic acid (MOPS), and 20 mM glucose, pH 7.0 with NaOH) supplemented with 2,3-butanedione monoxime (30 mM BDM; Sigma-Aldrich, St. Louis, MO, USA). The solutions were oxygenated with 100 % O2 at 37°C. After digestion with collagenase type I (288 U/ml; Worthington) and protease type XXIV (5 mg/ml; Sigma-Aldrich) for 15 min and agitation in protease-free solution for another 35 min, the cell suspension was filtered through a 200 nm mesh. Subsequently the Ca2+ concentration of the cell fraction was increased to 0.2 mM, the cell suspension was centrifuged and calcium tolerant rod-shaped single cardiomyocytes were declared as the cardiomyocyte fraction (CM). Remaining tissue chunks were declared as the non cardiomyocyte (NM) fraction. Cells were stored in TRIzol-Reagent (ThermoFisher) at −20°C. Isolation of total RNA was performed, using TRIzol-Reagent (ThermoFisher) and frozen tissue samples were processed according to the manufacturer’s protocol. Synthesis of single-stranded cDNA was carried out with the Maxima First Strand cDNA Synthesis Kit (ThermoFisher), using 3 μg of total RNA per 20 μl reaction. For quantitative real-time PCR (qPCR) 10 μl reactions, consisting of 0.5 μl cDNA, 5 μl TaqMan Fast Universal Master Mix (ThermoFisher), and 0.5 μl 6-carboxyfluorescein (FAM)-labeled TaqMan probes and primers (TaqMan Gene Expression Assays; ThermoFisher) were analyzed using the StepOnePlus (Applied Biosystems, Foster City, CA, USA) PCR system. hACE2 primers and probes (Hs01085333_m1) were purchased from ThermoFisher. The glyceraldehyde 3-phosphate dehydrogenase housekeeping gene (GAPDH: Hs99999905_m1) was used for normalization. All RT-qPCR reactions were performed as triplicates and control experiments in the absence of cDNA were included. Means of triplicates were used for the 2−ΔCt calculation, where 2−ΔCt corresponds to the ratio of mRNA expression versus GAPDH.

### Flow cytometry

Cells were dissociated to single-cell suspension with 0.25% trypsin (Gibco), then washed and incubated with Zombie-violet viability stain (Biolegend) at a dilution of 1:500 in 100 μL of PBS at room temperature. In some assays, antibodies against surface proteins were incubated with the cells. Fibroblasts were identified using anti-CD90 PE (Biolegend, catalog number 328110) and macrophages were identified using anti-CD14 APC (Biolegend, catalog number 367118). Cells then were fixed with 4% paraformaldehyde (final concentration) in phosphate-buffered saline for 20 min and analyzed on the MACSQuant 10 (Miltenyi Biotec) or the LSRFortessa X20 (Becton Dickinson Biosciences). Analysis was performed using FlowJo 10.6.1 (Becton Dickinson & Co).

### hPSC culture and cardiomyocyte differentiation

The parent stem cell line, BJFF.6, was generated from the human BJ fibroblast line (ATCC, catalog number: CRL-2522) by the Genome Engineering and iPSC Center at Washington University in St. Louis. This parent cell line has no known mutations associated with heart disease and the stem cells are pluripotent as assessed by immunofluorescence staining^56^. Differentiation to hPSC-derived CMs was performed as previously described ^56^. Briefly, stem cells were maintained in feeder-free culture and differentiation was initiated by temporal manipulation of WNT signaling^57,58^. hPSC-derived CMs were enriched using metabolic selection ^59^, yielding >90% cardiomyocytes ^56^. All experiments were conducted at least 30 days after the initiation of differentiation.

### Differentiation (and validation) of stem cell derived cardiac fibroblasts

hPSC derived cardiac fibroblasts were differentiated using the method of Zhang et al. (reference ^45^). Briefly, differentiation was initiated by directing the BJFF stem cell line towards a mesoderm/cardiac progenitor lineage by the activation and subsequent inhibition of WNT signaling using CHIR-99021 and IWP2 respectively. These cells were then directed to become proepicardial cells by the addition of retinoic acid and WNT CHIR 99021 followed by TGFβ inhibition (SB431542, Tocris Bioscience 1614). Finally, the pro-epicardial cells were directed to become quiescent cardiac fibroblasts by the addition of FGF2 and higher levels of TGFβ inhibition. Derivation of hPSC-cardiac fibroblasts was validated by measuring gene expression using RT-PCR. They showed high levels of both cardiac specific genes, GATA4 and TCF21 as well as general fibroblast genes COL1A1 and DDR2 (Fig. S1c).

### Fibroblast culture

BJ fibroblasts (ATCC, catalog number: CRL-2522) were cultured in Eagle’s Minimum Essential Medium with 10% FBS and 1% Pen-Strep (Gibco, catalog number: 15140122). Human ventricular cardiac fibroblasts that were harvested from normal adult ventricular tissue were obtained from Lonza, maintained at low passage number (<12), and cultured according to the manufacturer’s recommendations in FGM-3 growth media.

### Macrophage culture

CD34^+^ cells isolated from human cord blood were cultured in Iscove’s Modified Dulbecco’s Medium with 10% FBS, 1% Pen-Strep, and 10 ng/mL of human macrophage colony stimulating factor (M-CSF, R&D Systems, catalog number 216-MC) for at least 10 days to generate mature macrophages before use in experiments.

### Endothelial cell culture

H1 hPSCs were differentiated into vascular endothelial cells following the published protocol ^60^. Arterial endothelium was identified as CD34+CD184+CD73+ cells using appropriate antibodies (BD Biosciences, anti-CD34 PE-Cy7 [cat # 560710], anti-CD184 APC [cat #560936], anti-CD73 PE [cat #550257]) and isolated by flow cytometric cell sorting on a BD FACSAriaII. These cells were cultured in StemPro-34 serum free media (ThermoFisher, cat# 10639011).

### Isolation of primary human cardiac endothelial cells and macrophages

Human myocardium was dissected into ~200mg pieces and digested in 3mL DMEM containing 250U/mL collagenase IV, 60U/mL hyaluronidase and 60U/mL DNAseI for 45 minutes at 37 degrees C. Following digest and red blood cell lysis, the resultant single cell suspension was incubated with anti-CD14 PE (cat #301806), anti-CD64 PE-Cy7 (cat #305022), anti-CD45 FITC (cat #304006), and anti-CD31 BV421 (cat #303124) (all antibodies from Biolegend). Macrophages were identified as CD14+CD64+CD45+ cells and endothelial cells were identified as CD31+CD64-CD45− cells. Cells were isolated by flow cytometric cell sorting on a BD FACSMelody and cultured in StemPro-34 supplemented with either M-CSF or VEGF (R&D Systems, cat #293-VE).

### Two-dimensional cell cultures and tissues

hPSC-derived CMs, fibroblasts, and/or macrophages were dissociated from two-dimensional cultures using 0.25% Trypsin-EDTA, resuspended in media containing RPMI-1640 with 20% FBS and 10 μM Y-27632 and plated on gelatin coated tissue culture dishes. After 48 h, the media was changed to DMEM High glucose (4 mg/mL), 10% FBS, 1% non-essential amino acids, 1% GlutaMAX Supplement, and 1% Pen-Strep. All drug compounds were purchased from Selleckchem (ruxolitinib, catalog number S8932; MRT67307, catalog number S7948; E64, catalog number S7379; camostat, catalog number S2874) and resuspended to a stock concentration of 10μM in PBS or DMSO (depending on the solubility profile), then diluted to working concentration in culture media (described above) and sterile-filtered.

### Immunostaining of hPSC-derived CMs and confocal fluorescence microscopy

Immunostaining was performed as previously described with a few modifications^56^. Briefly, cardiomyocytes were fixed for 20 minutes in 4% formaldehyde in phosphate buffered saline (PBS). Cells were then permeabilized with 0.4% Triton X-100 for 20 minutes at room temperature. The cells were blocked for 1 hour using a blocking solution containing 3% bovine serum albumin, 5% donkey serum, 0.1% Triton X-100, and 0.02% sodium azide in PBS. Primary antibodies (rabbit anti Troponin T, 1:400, Abcam, ab45932) were added for 1-2 hours at room temperature or overnight at 4 °C. Cells were then washed with PBS before incubating for 1 hour in secondary antibody (Cy3 donkey anti-rabbit, Jackson Immunoresearch, 711165152). 4′,6-diamidino-2-phenylindole (DAPI) was used at a 1:50000 dilution to stain for nuclei. Cells were visualized using a Nikon A1Rsi confocal microscope (Washington University Center for Cellular Imaging). Z-stacks of cells with 40x magnification were recorded in sequential scanning mode. Images were processed in ImageJ and Z-stacks were converted to standard deviation projections^61^.

### Engineered heart tissues (EHTs)

EHTs were prepared according to published protocol^62^ with modifications. Casting molds for tissue formation were prepared using PDMS at a 1:25 ratio. 1 mL of PDMS was poured into each well of a 24-well plate and a Teflon spacer (EHT Technologies GmBH; Hamburg, Germany) was placed inside to generate a well. The PDMS was degassed under high vacuum and baked overnight at 65°C. The Teflon spacers were removed using ethanol.

Prior to use, the casting molds were sterilized with ethanol, dried with nitrogen gas, and placed under UV light for 10-15 min. 1% pluronic-F127 in PBS was added to the molds for 20 min to block the surface from adhering to the seeded tissues. The pluronic was removed, the casting molds were rinsed twice with PBS, and then dried. Silicone racks consisting of two pairs of PDMS posts (EHT Technologies GmBH, Hamburg, Germany) were positioned such that each pair of PDMS posts fit within one casting mold.

The procedure for seeding of tissues was modified from^62^. Working on ice, rat collagen I (1 mg/mL final concentration) was combined with equal parts 2x DMEM containing FBS and neutralized with sodium hydroxide. Growth factor reduced Matrigel (Corning, catalog number: 354230) was added to a final concentration of 0.77 g/mL. hPSC-derived CMs, fibroblasts, and/or macrophages were dissociated from two-dimensional cultures with 0.25% Trypsin-EDTA and the Trypsin was quenched in RPMI-1640 with 20% FBS media containing 10 g/mL of DNaseI. Cells were then centrifuged, resuspended in media containing RPMI-1640 with 20% FBS and 10 μM Y-27632, and combined with the collagen/Matrigel mixture. Each tissue contained 10^6^ hPSC-derived CMs, 5% fibroblasts, and 10% macrophages. Tissues were seeded in the casting molds with the silicone racks in a 100 μL volume. After polymerizing around the silicone racks for 2 h at 37°C, the tissues were covered overnight with RPMI-1640 containing 20% FBS. The tissues, attached to the posts of the silicone racks, then were moved out of the casting molds and transferred into media containing DMEM High glucose (4 mg/mL), 10% FBS, 1% non-essential amino acids, 1% GlutaMAX Supplement, and 1% Pen-Strep. Consistent with previous reports, engineered heart tissue contraction was observed ~2-5 days after seeding, and the displacement increased over time as the tissues organize and mature^62^. By day 7 post seeding, all tissues generated aey t least 0.025 mm of displacement (see details below). EHTs were inoculated with SARS-CoV-2 at least 7 days after tissue seeding, and EHT contraction and morphology were measured daily throughout the course of the experiment.

### Analysis of engineered heart tissue contractility

EHTs between two PDMS posts were visualized on an EVOS microscope and videos of spontaneously contracting posts were recorded at 30 frames per second using a Macintosh desktop with built-in camera (**Supplementary Videos 1 and 2**). We used the 2 mm diameter of the caps on the posts to calibrate the pixels per mm for each video. We wrote a custom script in MATLAB to calculate the displacement of the posts as a function of time. Automated tracking was done using the computer vision toolbox and the displacement was calculated as a function of time. A second order polynomial spline fit was applied to remove any drift in the camera position. Traces were smoothed using a Savitsky-Golay filter and peaks in the displacement were identified using the findpeaks algorithm. The average and standard deviation of the displacement was then calculated for each ~60 sec video. The time for force development was defined as the time required to achieve 75% activation and the time for relaxation was defined as the time to relax to 75% of the peak activation.

### Histology of autopsy and engineered heart tissues

Tissues were fixed with 10% NBF for 7 days, embedded in 1% agar, mounted in cassettes, and embedded in paraffin. Target markers were visualized using Opal 4-Color Manual IHC Kit (Perkin-Elmer) with the following changes: 1) 10% NBF fixation step was substituted for treatment with 10% MeOH + 10% hydrogen peroxide in water for 20 min; 2) during antigen retrieval, the vessel with AR6 was brought to boiling and slides were immediately transferred into deionized water; 3) blocking buffer was substituted for 10% FBS in TBST. Primary antibodies against human sarcomeric actin (Sigma A2127), human Troponin T (ThermoFisher MS-295-P1), human CD68 (BioRad MCA5709), human ACE2 (Abcam ab15348), human Ki67 (Abcam ab16667), human CCR2 (Abcam ab176390), SARS-CoV-2-N (Sino Biological 40143-R001) were used. Cell death was assessed using TUNEL staining from *In Situ* Cell Death Detection Kit (Roche) with Opal-based costain for sarcomeric actin as described above. Viral RNA was directly visualized with RNAscope Multiplex Fluorescent Reagent Kit v2 Assay (Advanced Cell Diagnostics) and RNAscope 2.5 HD Detection Reagent - RED (Advanced Cell Diagnostics) using positive-strand and negative strand probes for ORF1ab. Images were collected on a confocal microscope (Zeiss LSM 700 Laser Scanning Confocal) and processed using ZenBlack (Zeiss) and ImageJ (NCBI). Troponin staining was quantified using manual cell tracing in ImageJ (NCBI) from at least three areas and 30 cells analyzed per tissue.

### Electron microscopy

Cells grown on aclar coverslips were briefly rinsed in 0.15 M cacodylate buffer that was warmed to 37°C followed by the addition of 2.5% glutaraldehyde, 2% paraformaldehyde, 0.2% tannic acid in 0.15 M cacodylate buffer with 2 mM CaCl_2_, pH 7.4 at 37°C. Once added, the coverslips were returned to a 37°C incubator for 15 min followed by overnight fixation at room temperature. Post fixation, samples were rinsed in 0.15 M cacodylate buffer 4 times for 15 min each followed by a secondary fixation in 1% OsO_4_/1.5% K_3_Fe(CN)_6_ in 0.15 M cacodylate buffer for 1.5 h in the dark. The coverslips were then rinsed 4 times in ultrapure water for 15 min each followed by *en bloc* staining with 2% aqueous uranyl acetate overnight at 4°C in the dark. After another 4 water washes, the samples were dehydrated in a graded ethanol series (30%, 50%, 70%, 90%, 100% x4) for 10 min for each step. Once dehydrated, cells were infiltrated with LX112 resin over a period of 2 days. The coverslips then were flat embedded and polymerized at 60°C for 48 h. Once polymerized, the aclar coverslips were peeled away from the resin, and small areas were excised and mounted perpendicularly on a blank epoxy stub for cross sectioning. 70 nm sections were then cut and imaged on a TEM (JEOL JEM-1400 Plus) at 120 KeV.

### RNA sequencing and analysis

Samples were prepared according to library kit manufacturer’s protocol, indexed, pooled, and sequenced on an Illumina NovaSeq 6000. Basecalls and demultiplexing were performed with Illumina’s bcl2fastq software and a custom python demultiplexing program with a maximum of one mismatch in the indexing read. RNA-seq reads were then aligned to the Human Ensembl GRCh38.76 primary assembly and SARS-CoV-2 NCBI NC_045512 Wuhan-Hu-1 genome with STAR version 2.5.1a^63^. Gene counts were derived from the number of uniquely aligned unambiguous reads by Subread:featureCount version 1.4.6-p5^64^. Isoform expression of known Ensembl transcripts were estimated with Salmon version 0.8.2^65^. Sequencing performance was assessed for the total number of aligned reads, total number of uniquely aligned reads, and features detected. The ribosomal fraction, known junction saturation, and read distribution over known gene models were quantified with RSeQC version 2.6.2^66^.

All gene counts were imported into the R/Bioconductor package EdgeR^67^ and TMM normalization size factors were calculated to adjust for samples for differences in library size. Ribosomal genes and genes not expressed in at least four samples greater than one count-per-million were excluded from further analysis. The TMM size factors and the matrix of counts were then imported into the R/Bioconductor package Limma^68^. Weighted likelihoods based on the observed mean-variance relationship of every gene and sample were then calculated for all samples with the voomWithQualityWeights^69^. The performance of all genes was assessed with plots of the residual standard deviation of every gene to their average log-count with a robustly fitted trend line of the residuals. Differential expression analysis was then performed to analyze for differences between conditions and the results were filtered for only those genes with Benjamini-Hochberg false-discovery rate adjusted p-values less than or equal to 0.05.

For each contrast extracted with Limma, global perturbations in known Gene Ontology (GO) terms, MSigDb, and KEGG pathways were detected using the R/Bioconductor package GAGE^70^ to test for changes in expression of the reported log 2 fold-changes reported by Limma in each term versus the background log 2 fold-changes of all genes found outside the respective term. The R/Bioconductor package heatmap3^71^ was used to display heatmaps across groups of samples for each GO or MSigDb term with a Benjamini-Hochberg false-discovery rate adjusted p-value less than or equal to 0.05. Perturbed KEGG pathways where the observed log 2 fold-changes of genes within the term were significantly perturbed in a single-direction versus background or in any direction compared to other genes within a given term with p-values less than or equal to 0.05 were rendered as nnotated KEGG graphs with the R/Bioconductor package Pathview^72^.

To find the most critical genes, the raw counts were variance stabilized with the R/Bioconductor package DESeq2^54^ and then analyzed via weighted gene correlation network analysis with the R/Bioconductor package WGCNA^73^. Briefly, all genes were correlated across each other by Pearson correlations and clustered by expression similarity into unsigned modules using a power threshold empirically determined from the data. An eigengene was created for each de novo cluster and its expression profile was then correlated across all coefficients of the model matrix. Because these clusters of genes were created by expression profile rather than known functional similarity, the clustered modules were given the names of random colors where grey is the only module that has any pre-existing definition of containing genes that do not cluster well with others. These *de novo* clustered genes were then tested for functional enrichment of known GO terms with hypergeometric tests available in the R/Bioconductor package clusterProfiler^74^. Significant terms with Benjamini-Hochberg adjusted p-values less than 0.05 were then collapsed by similarity into clusterProfiler category network plots to display the most significant terms for each module of hub genes in order to interpolate the function of each significant module. The information for all clustered genes for each module were combined with their respective statistical significance results from Limma to identify differentially expressed genes.

### Statistical analysis

Statistical tests were chosen based on standards in the virology and cardiomyocyte fields for given assay. Parametric and non-parametric statistical methods were used when appropriate. Statistical significance was assigned when *P* values were < 0.05 using Prism Version 8 (GraphPad). Specific tests are indicated in the figure legends.

## Data availability

All data supporting the findings of this study are found within the paper and its Extended Data Figures and are available from the corresponding authors upon request. RNA sequencing data sets generated in this study will be made available in GEO at the time of publication.

## Acknowledgements

The authors would like to acknowledge funding support from the National Institutes of Health (R01HL141086 to M.J.G, R01 HL138466 to K.J.L., R01 HL139714 to K.J.L., 75N93019C00062 and R01 AI127828 to M.S.D.), Burroughs Welcome Fund (1014782 to K.J.L.), Defense Advanced Research Project Agency (HR001117S0019), the March of Dimes Foundation (FY18-BOC-430198 to M.J.G.), Foundation of Barnes-Jewish Hospital (8038–88 to K.J.L.), and Children’s Discovery Institute of Washington University and St. Louis Children’s Hospital (CH-II-2017–628 to K.J.L., PM-LI-2019-829 to K.J.L. and M.J.G.). Imaging was performed in the Washington University Center for Cellular Imaging (WUCCI) which is funded, in part by the Children’s Discovery Institute of Washington University and St. Louis Children’s Hospital (CDI-CORE-2015-505 and CDI-CORE-2019-813) and the Foundation for Barnes-Jewish Hospital (3770). The authors thank Dr. Cynthia Goldsmith for help interpreting electron microscopy micrographs and the McDonnell Genome Institute (MGI) at Washington University School of Medicine for assistance in performing sequencing and analysis.

## Author Contributions

Conceptualization, M.J.G, A.L.Bailey, K.L, M.S.D.; Methodology, L.G., W.T.S., A.L.Bailey; Software, M.J.G.; Investigation, L.G., O.D., A.L.Bailey, A.L.Bredemeyer, P.M., J.L., V.P., E.W., S.S, J.A.F., W.G., C.S.; Resources, E.B., A.J.N, K.A.H., A.S.R, M.S., D.H., X.X., P-Y.S, J.T.H, F.L., W.T.S., L.S.; Writing – Original Draft, L.G., M.J.G., A.L.Bailey, K.L., O.D., A.L.Bredemeyer; Writing – Review & Editing, L.G., M.J.G., A.L.Bailey, K.L., O.D., A.L.Bredemeyer, M.S.D.; Funding Acquisition, M.J.G, K.L., M.S.D; Supervision, M.S.D., M.J.G, C-Y.L., K.L.

## Key Resources Table

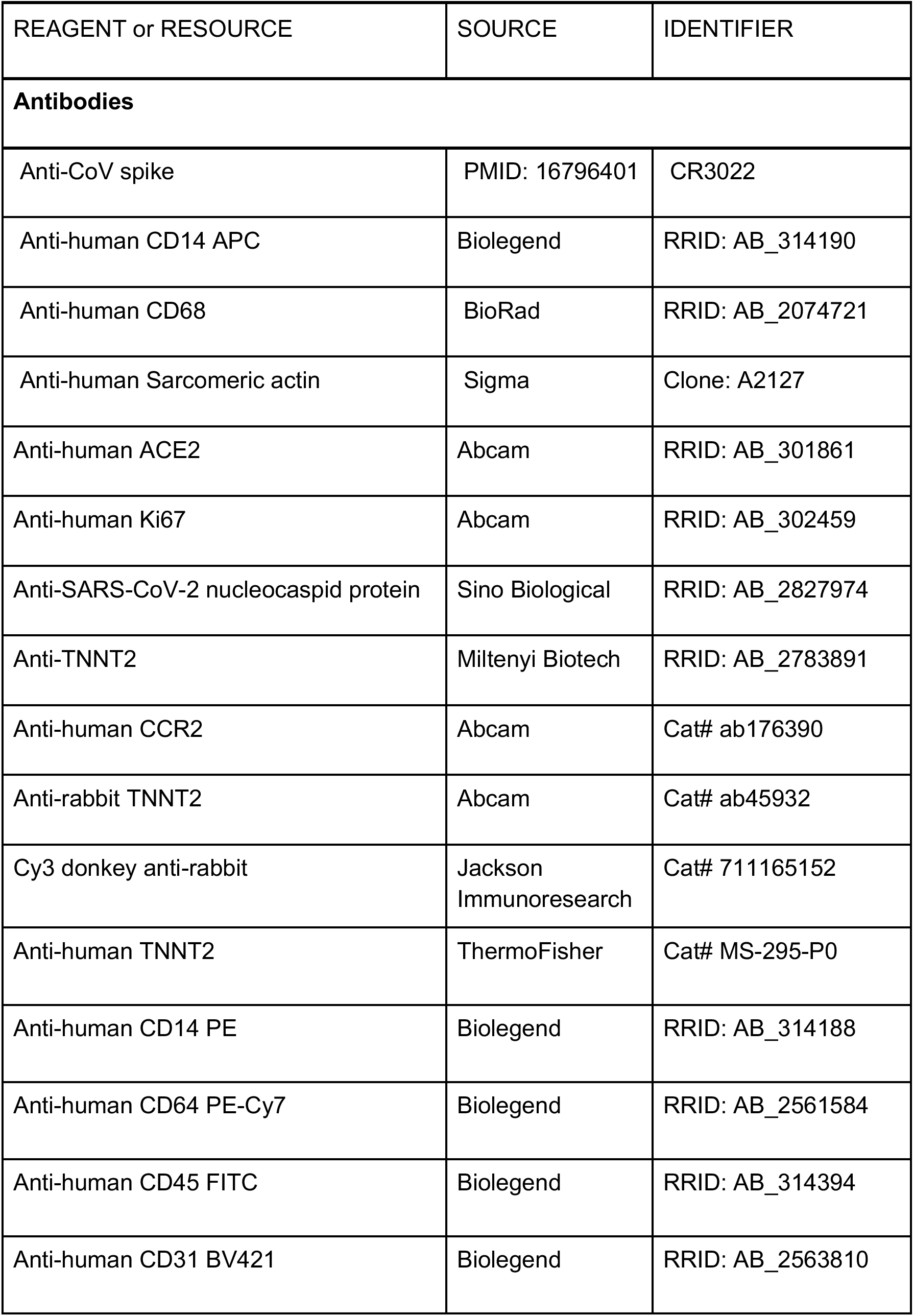
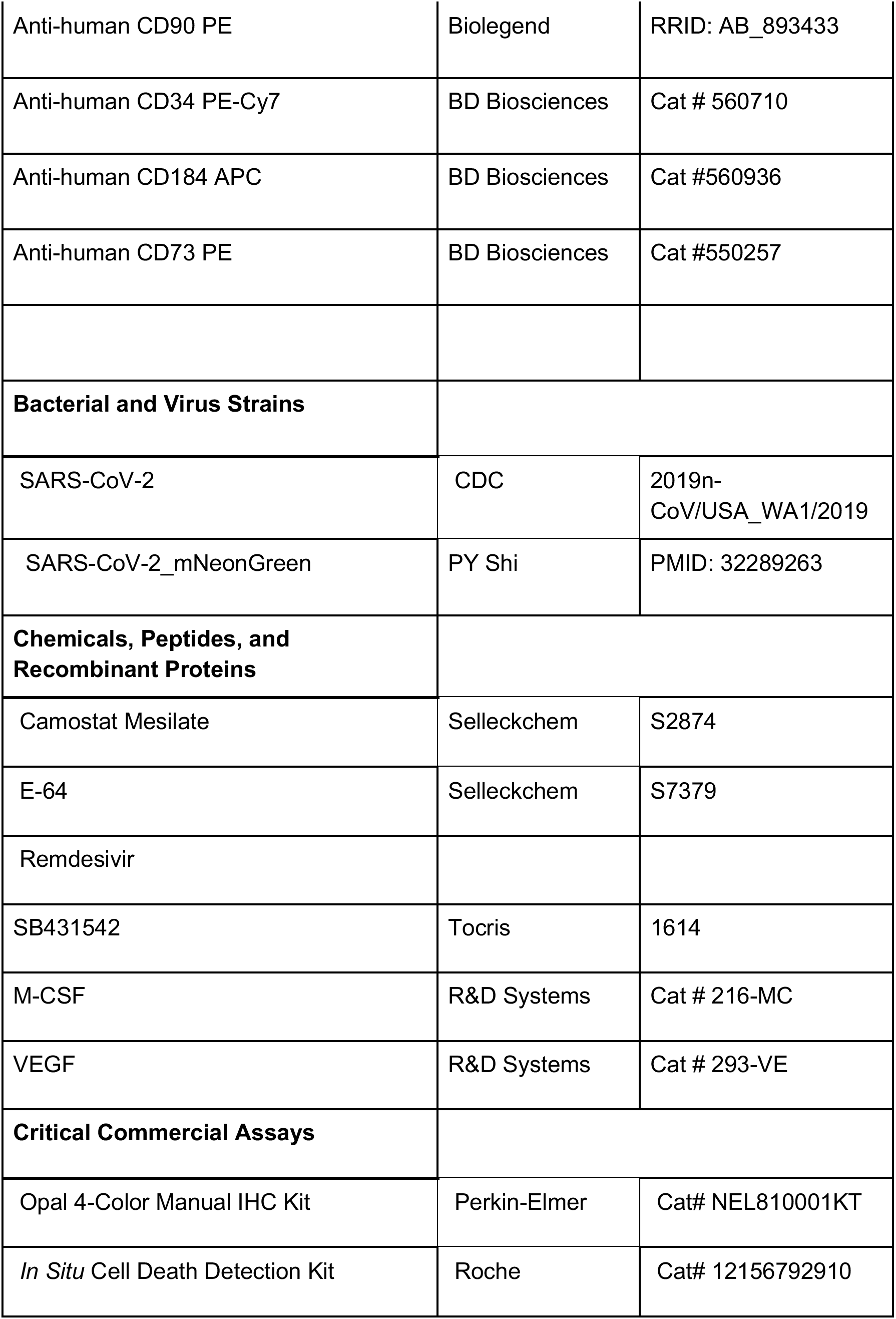
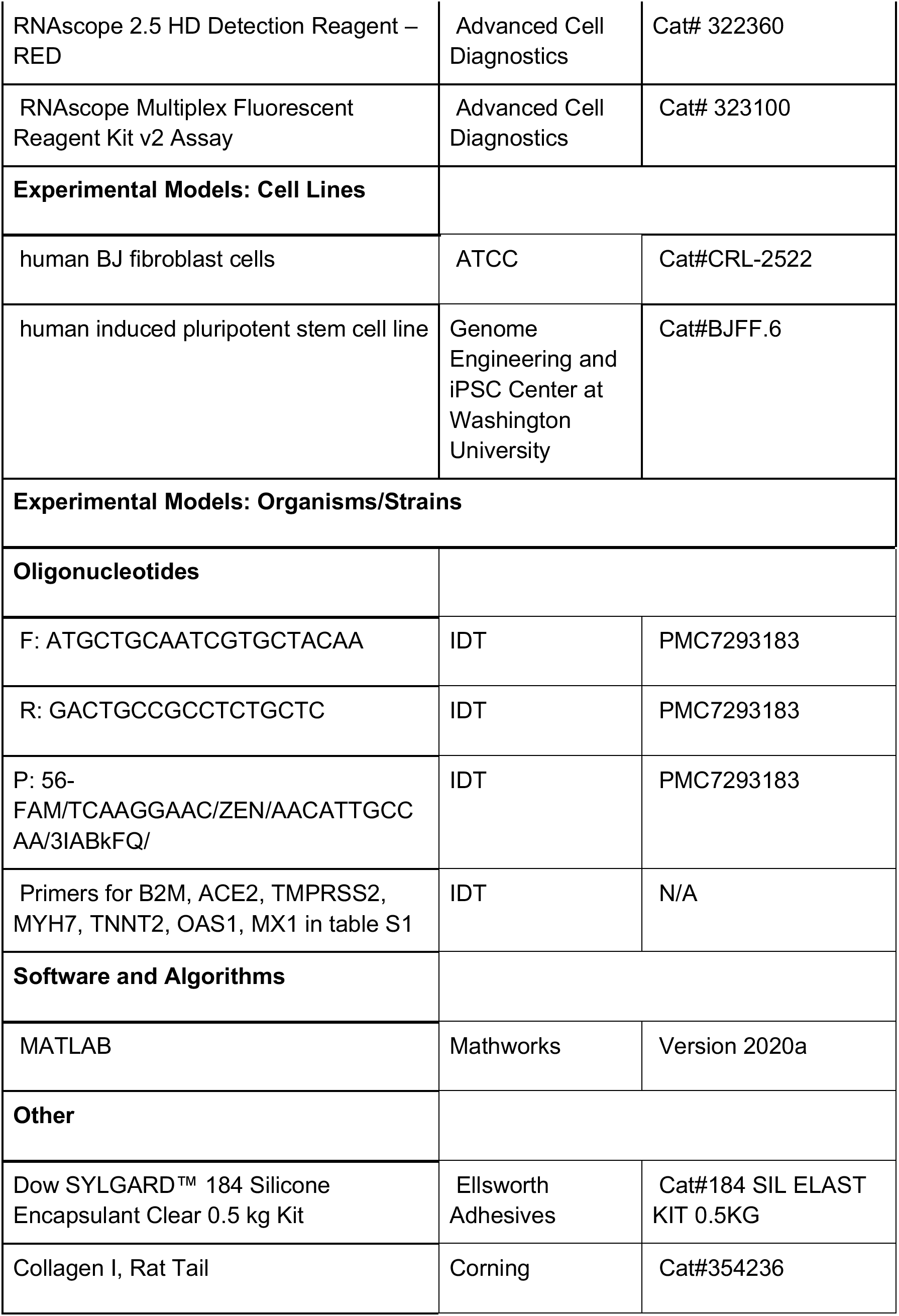
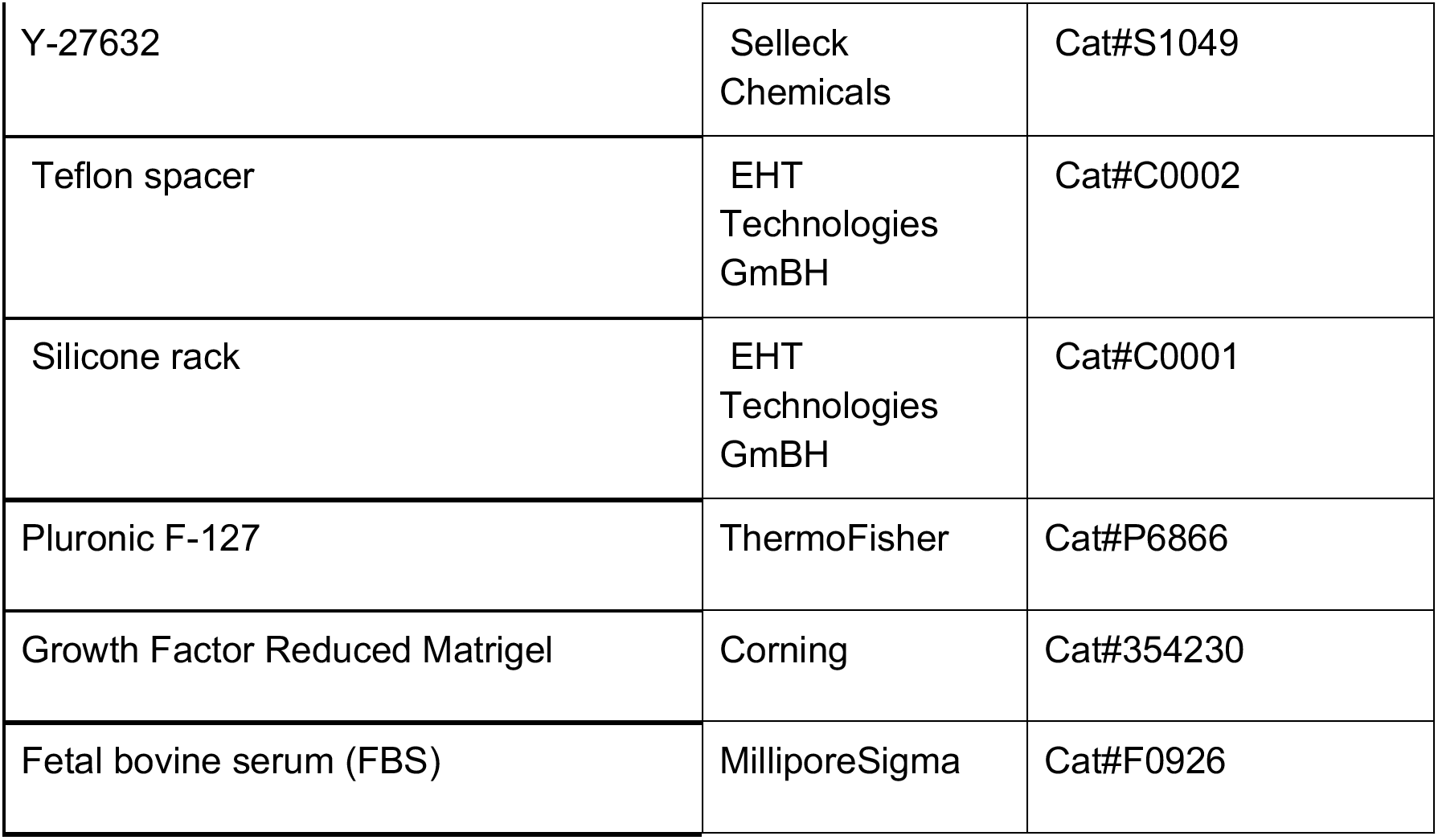

**Table S1.**
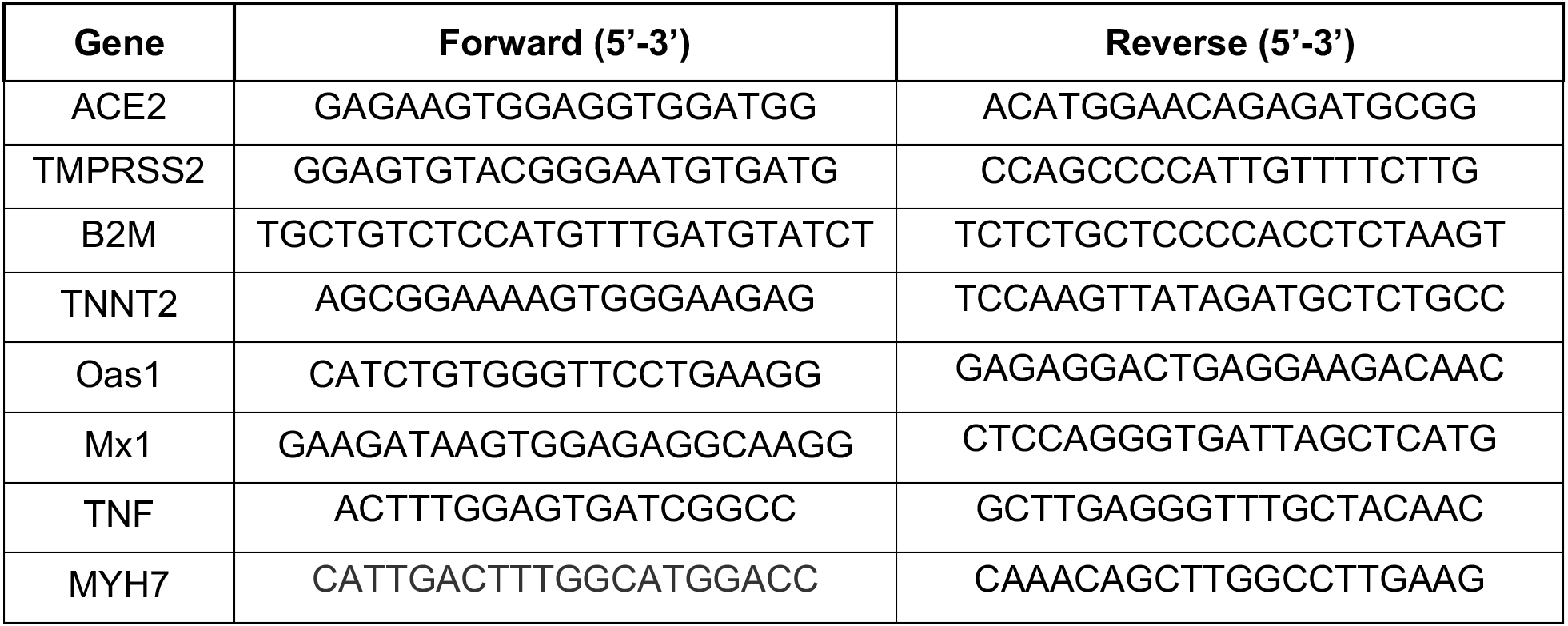
Primers for qPCR-base evaluation of human host gene expression

