## Supplemental Figures for "SARS-CoV-2 Infects Human Engineered Heart Tissues and Models COVID-19 Myocarditis"

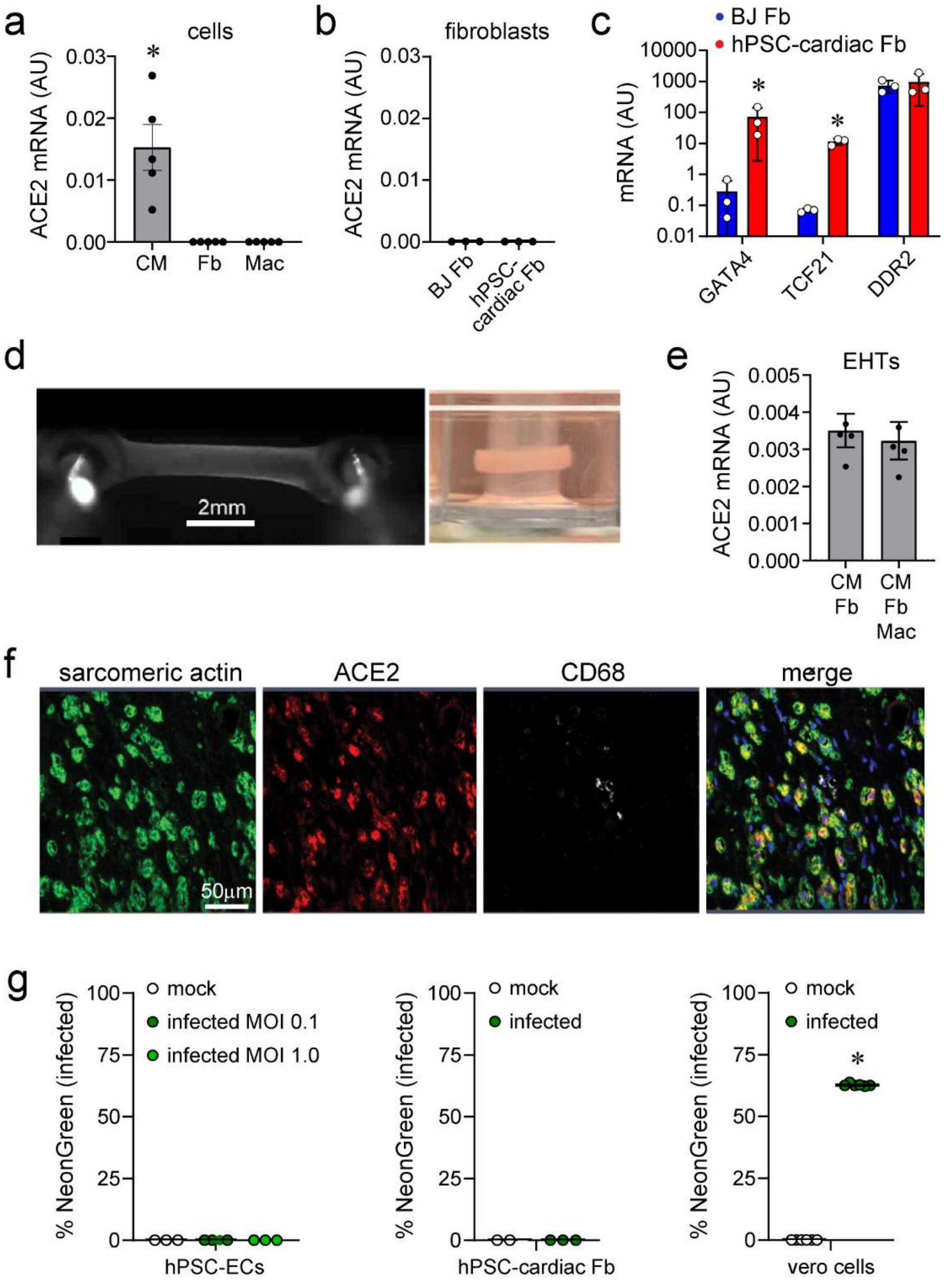


**Figure S1. ACE2 expression and infectivity of hPSC-derived cells in human engineered heart tissues. a,** Quantitative RT-PCR measurements showing *ACE2* mRNA expression in hPSC-derived cardiomyocytes (CM). *ACE2* was not detected in human fetal cord blood-derived macrophages (Mac) or dermal fibroblasts (Fb). Each data point represents an independent experiment (n=5) and error bars denote standard error of the mean. **b**, *ACE2* was not detectable in hPSC-derived cardiac fibroblasts or BJ dermal fibroblasts. Each data point represents an independent experiment (n=3). **c**, Quantitative RT-PCR showing marked enrichment of GATA4 *and* TCF21 mRNA expression in hPSC-derived cardiac fibroblasts compared to BJ fibroblasts.

Both fibroblast populations express DDR2. Data is presented on a log10 scale. Each data point represents an independent experiment (n=5), bar height corresponds to the mean value, and error bars denote standard deviation. * denotes $p<0.05$ compared to BJ fibroblasts (Mann-Whitney test). **d,** Human engineered heart tissue formed between two deformable PDMS posts. When the tissue contracts, it displaces the posts (left). Ring shaped human engineered heart tissue formed on a single PDMS post (right). **e**, Quantitative RT-PCR measurements of *ACE2* mRNA expression in engineered heart tissues (EHTs). Data points indicate individual samples (n=4). Bars denote the mean value and error bars reflect standard error of the mean. * denotes $p<0.05$ compared to other groups (Mann-Whitney test). **f**, Immunohistochemistry of EHTs demonstrating ACE2 expression (red) in hPSC-derived cardiomyocytes (green, sarcomeric actin). Macrophages (white, CD68). DAPI: blue. Represented image from 7 analyzed specimens. **g**, Inoculation of hPSC-derived endothelial cells (EC) and hPSC-derived cardiac fibroblasts with mock (black) or SARS2-CoV-2-NeonGreen (green, MOI 0.1, 1.0). Vero cells are included as a positive control. Each data point represents biological replicates and bars denote mean values.. * $p<0.05$ compared to mock infection (Mann-Whitney test).

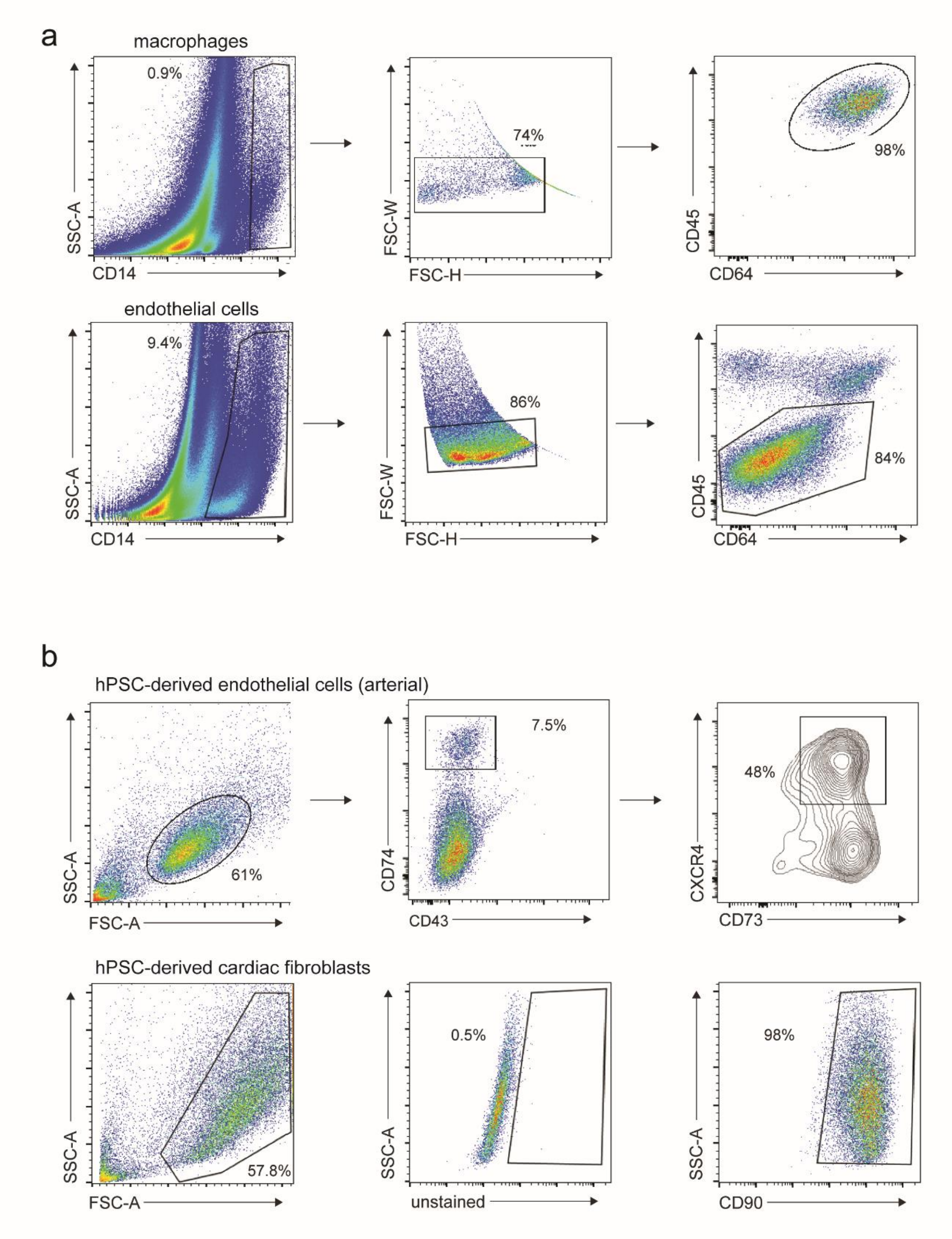


**Figure S2. Flow cytometry gating strategies for primary human cardiac cell types and hPSC-derived endothelial cells and cardiac fibroblasts. a**, Flow cytometry gating scheme for primary cardiac macrophages and endothelial cells. **b**, Flow cytometry gating strategies for hPSC-derived endothelial cells and cardiac fibroblasts. Representative plots and percentages are shown.

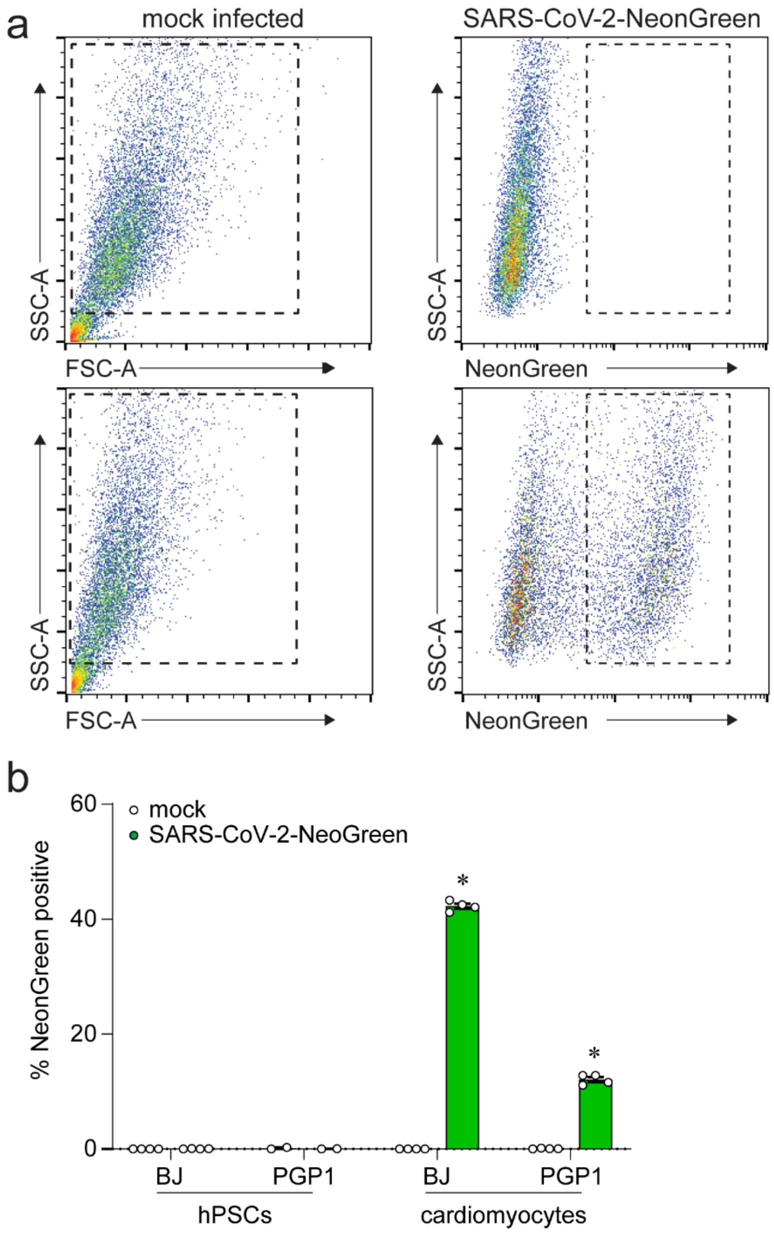


**Figure S3. SARS-CoV-2 infects hPSC-derived cardiomyocytes but not undifferentiated hPSC lines. a**, Flow cytometry gating scheme. hPSC-derived cardiomyocytes were either mock infected or inoculated with SARS-CoV-2-mNeonGreen (MOI 0.1) and harvested 3 days later. NeonGreen expression was measured using flow cytometry. **b**, Two independent undifferentiated hPSCs (BJ and PGP1 cell lines) or hPSC-derived cardiomyocytes were mock-infected or inoculated with SARS-CoV-2-mNeonGreen (MOI 0.1) and harvested 3 days later. The percentage of NeonGreen positive cells was quantified by flow cytometry. Data points indicate individual samples. Error bars denote standard deviation and bars correspond to the mean value. * $p<0.05$ compared to mock infection (Mann-Whitney test).

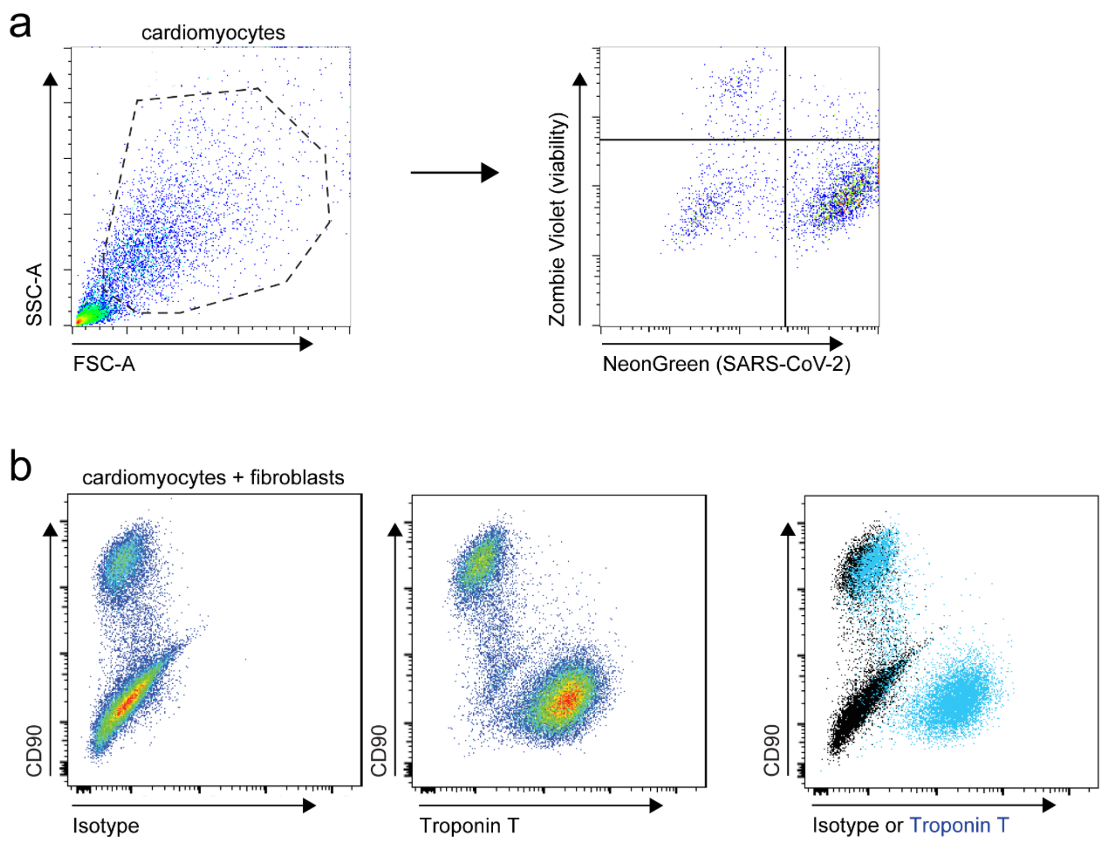


**Figure S4. Flow cytometry gating schemes. a,** hPSC-derived cardiomyocytes were infected with SARS-CoV-2-NeonGreen (MOI 0.1) and harvested 4 days after inoculation. Cells were stained with Zombie-Violet viability dye. Representative plot from 4 independent experiments. **b**, Mixed two-dimensional cultures of hPSC-derived cardiomyocytes and fibroblasts were permeabilized and stained with anti-CD90 and anti-Troponin T (TNNT2) antibodies to confirm that the CD90- cells in **Fig. 2f** are cardiomyocytes (CD90-TNNT2+).

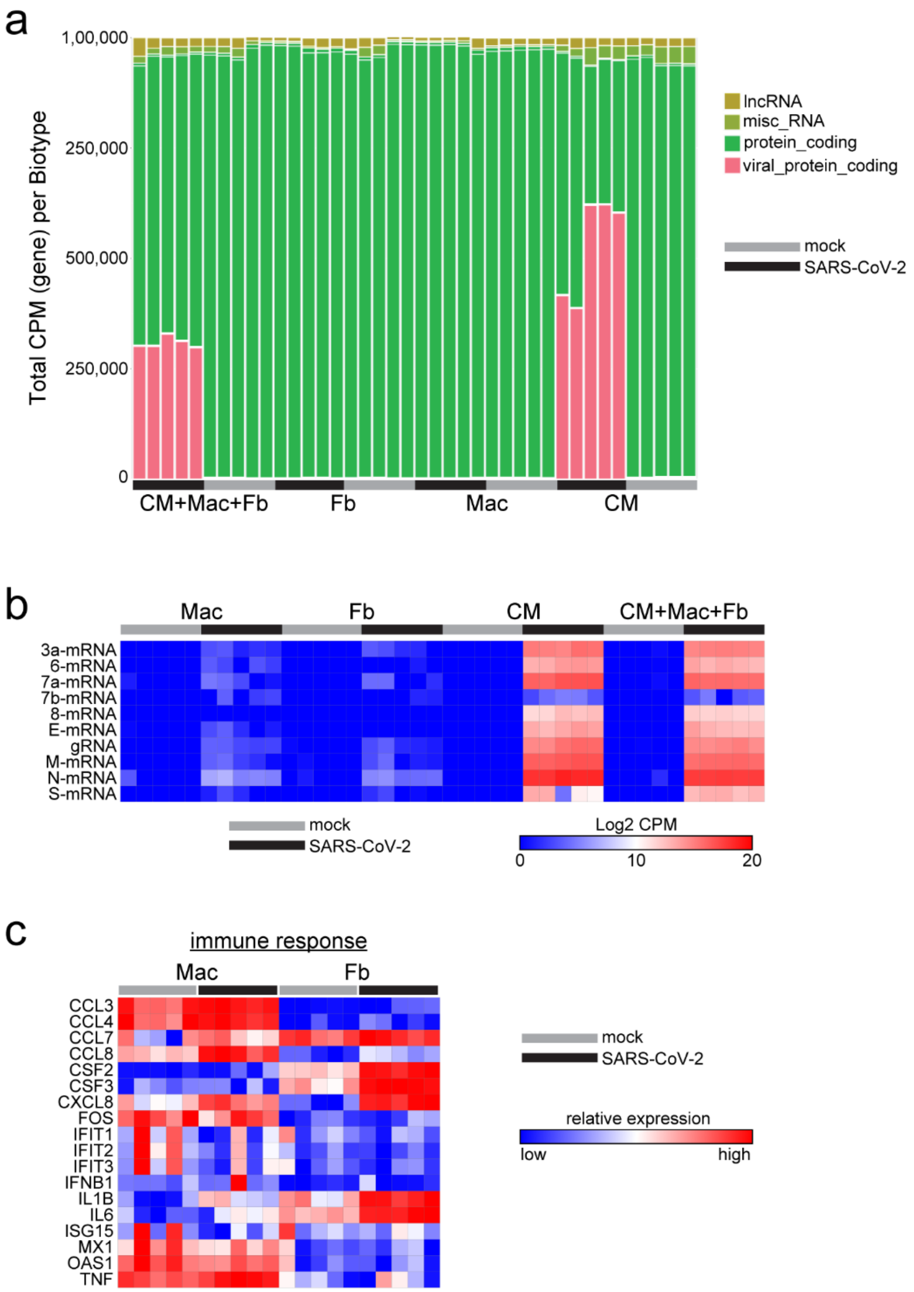


**Figure S5. RNA sequencing of mock and SARS-CoV-2 infected hPSC-derived cardiomyocytes, fibroblasts, macrophages, and two-dimensional tissues. a**, Biotype plot showing different classes of RNAs identified in RNA sequencing of mock (grey) and SARS-CoV-2 (black) infected cells. CPM: counts per million. **b**, Heat map showing the absolute expression of viral subgenomic RNAs. **c**, Heat map showing relative expression of chemokines, cytokines, and interferon stimulated transcripts in mock and SARS-CoV-2 infected macrophages and fibroblasts. Colors correspond to the relative expression of each transcript across cell types and experimental conditions. CM: hPSC-derived cardiomyocyte, Fb: fibroblast, Mac: macrophage.

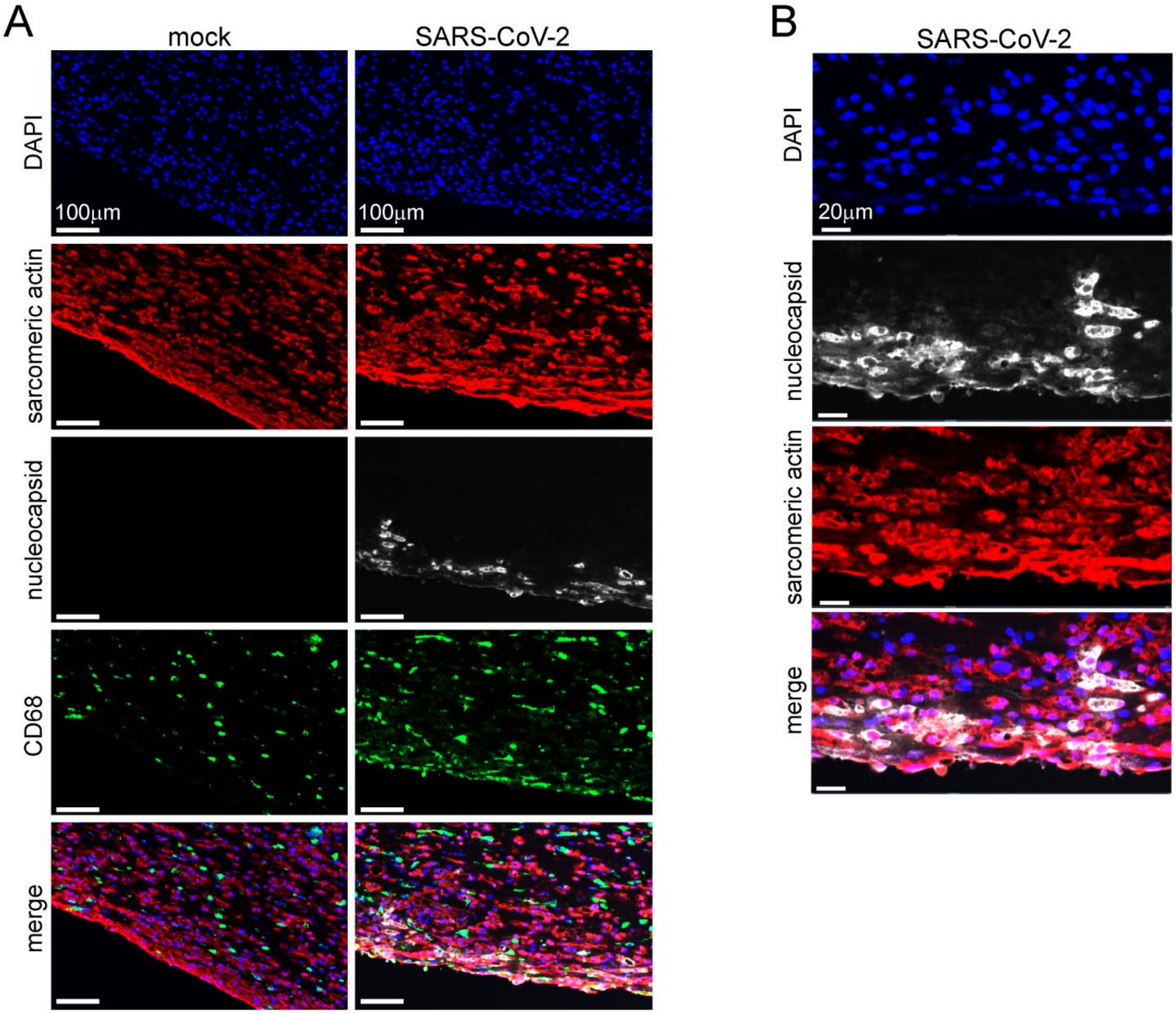


**Figure S6. Confocal microscopy of mock and SARS-CoV-2 infected three-dimensional EHTs. a**, Images of mock and SARS-CoV-2 infected three-dimensional EHTs stained with sarcomeric actin (cardiomyocytes, red), nucleocapsid protein (white), and CD68 (macrophages, green). **b**, High magnification images of SARS-CoV-2 infected three-dimensional EHTs demonstrating co-localization of nucleocapsid (white) and sarcomeric actin (red) staining. EHTs were harvested 5 days after infection. Images are representative of 4 independent experiments.

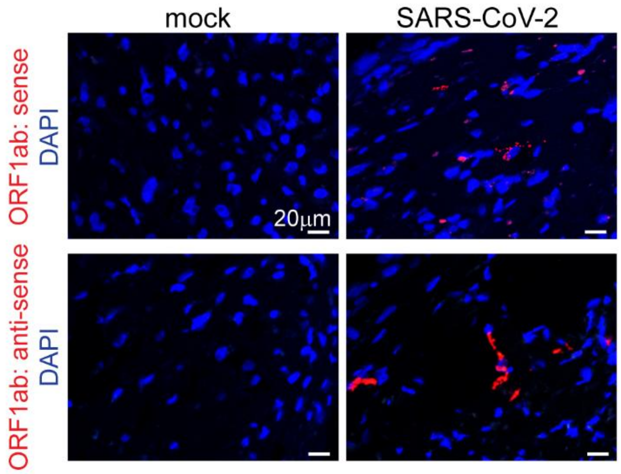


**Figure S7. RNA *in situ* hybridization for SARS-CoV-2 in EHTs.** *In situ* hybridization for SARS-CoV-2 ORF1ab (spike) RNA sense and anti-sense strands (red) in EHTs 5 days after mock or SARS-CoV-2 infection (MOI 0.1). DAPI: blue. Representative images from 4 independent specimens.

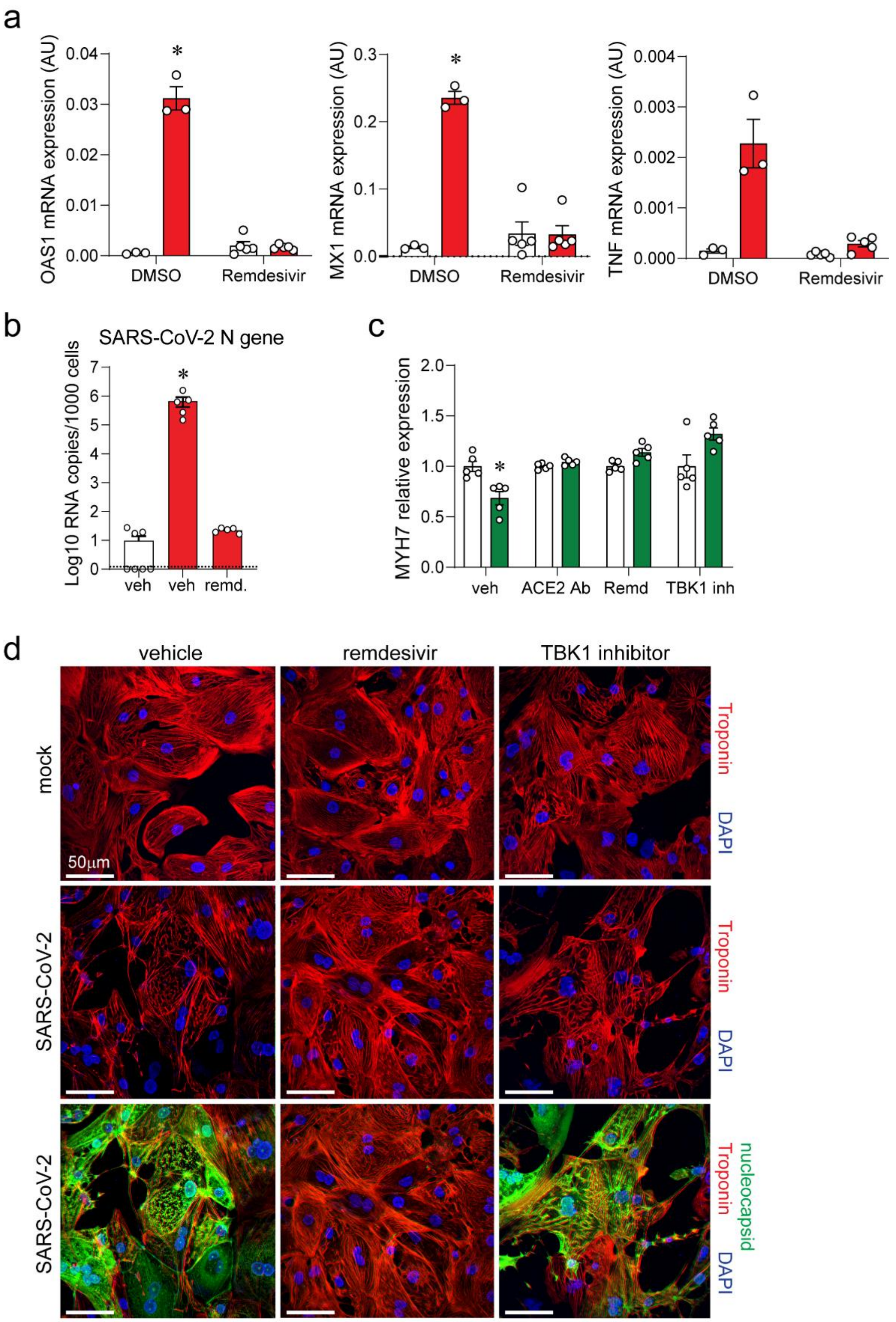
a
OAS1 mRNA expression (AU)
MX1 mRNA expression (AU)
TNF mRNA expression (AU)
DMSO
Remdesivir
b
SARS-CoV-2 N gene
Log10 RNA copies/1000 cells
veh
veh
remd.
c
MYH7 relative expression
veh
ACE2 Ab
Remd
TBK1 inh
d
vehicle
remdesivir
TBK1 inhibitor
mock
SARS-CoV-2
SARS-CoV-2
Troponin
DAPI
nucleocapsid
50μm

**Figure S8. SARS-CoV-2 infection drives type I interferon signaling and sarcomere breakdown in hPSC-derived cardiomyocytes. a,** Quantitative RT-PCR measuring OAS1, MX1, and TNF mRNA expression in 3D EHTs 5 days after inoculation with mock control (white) or SARS-CoV-2 (red, MOI 0.1). EHTs were treated with vehicle (DMSO) or remdesivir (10μM). Each data point denotes a biologically unique sample, bar height corresponds to the mean, and error bars indicate standard error of the mean. * $p<0.05$ compared to vehicle control. **b,** EHTs were either mock infected (white) or inoculated with SARS-CoV-2 (red, MOI 0.1). EHTs were treated with either vehicle control, 10 μM or 10 μM of remdesivir (remd.). Five days post-infection EHTs were homogenized and SARS-CoV-2 N gene RNA was measured by quantitative RT-PCR. Each data point denotes an individual EHT, bar height corresponds to the mean, error bars represent standard error of the mean, dotted line denotes lower limit of detection, *$p<0.05$ compared to mock vehicle (Mann-Whitney test). **c,** Quantitative RT-PCR measuring MYH7 mRNA expression in hPSC-derived cardiomyocytes 3 days after inoculation with mock control (white) or SARS-CoV-2 (green, MOI 0.1). Cells were treated with vehicle, ACE2 Ab (20μg/ml), remdesivir (10μM), or TBK inhibitor (MRT67307, 10μM). Each data point denotes a biologically unique sample, bar height corresponds to the mean, and error bars indicate standard error of the mean. * $p<0.05$ compared to mock control. **d**, Immunostaining of hPSC-derived cardiomyocytes for Troponin T (red) 3 days after inoculation with mock control or SARS-CoV-2-mNeonGreen (MOI 0.1). hPSC-derived cardiomyocytes were treated with vehicle, remdesivir (10μM) or TBK inhibitor (MRT67307, 10μM). Endogenous NeonGreen florescence is shown (nucleocapsid, green). Blue: DAPI.

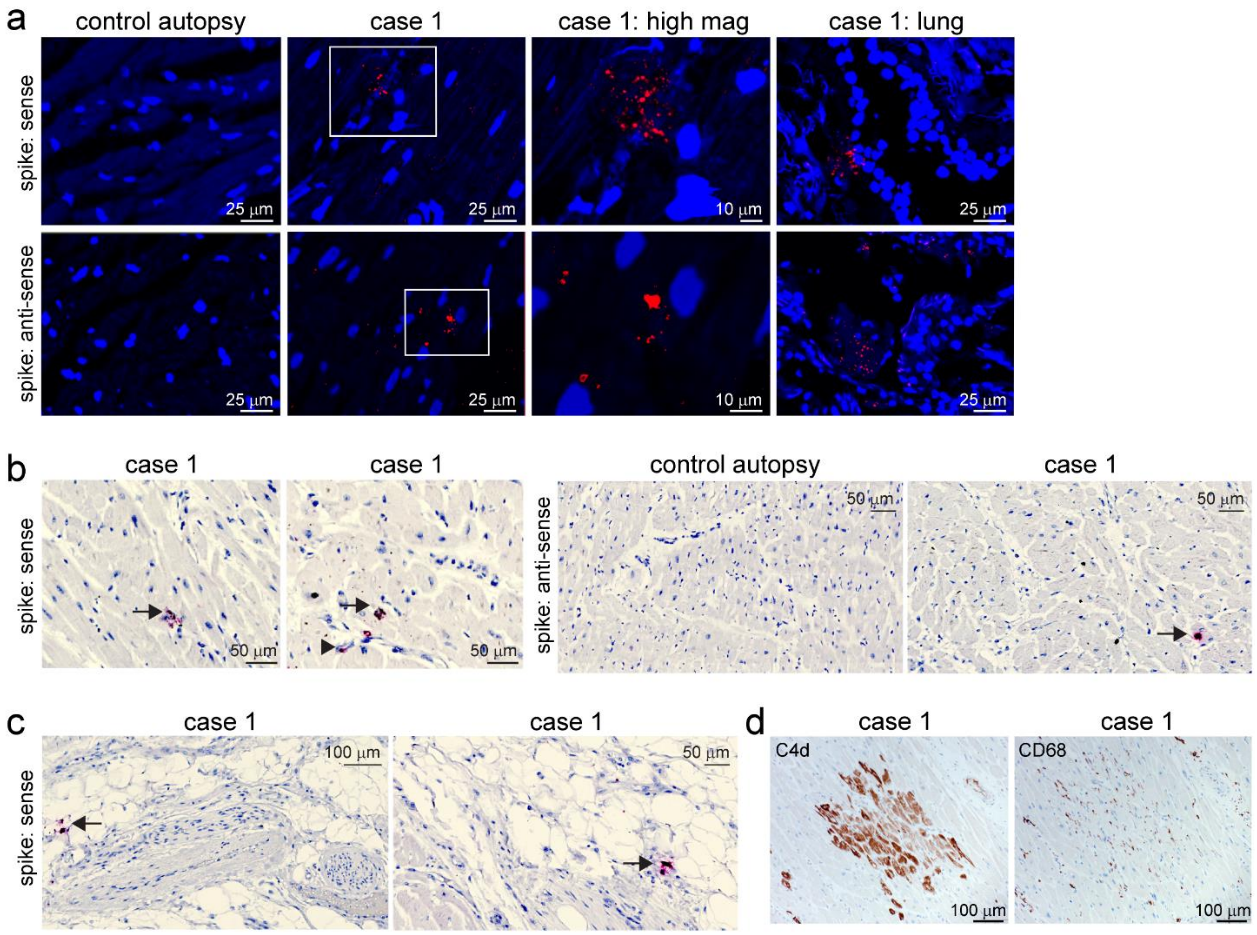


**Figure S9. I*n situ* hybridization for SARS-CoV-2 RNA in cardiac autopsy specimens. a,** *in situ* hybridization of cardiac autopsy and lung autopsy tissue for SARS-CoV-2 spike RNA showing evidence of viral infection. Red: spike RNA probes, Blue: DAPI. **b**, *in situ* hybridization of cardiac autopsy for SARS-CoV-2 (red) in control and COVID-19 myocarditis autopsy tissue. Staining is present within cardiomyocytes found in the COVID-19 myocarditis specimen (arrow). Arrowhead indicates spike staining in perivascular cells. No staining was evident in the control cases. Blue: hematoxylin. **c**, Spike sense probe staining (red) in epicardial adipose tissue. Staining (arrows) is present in perivascular adipocyte tissue. Blue: hematoxylin. **d,** Immunostaining of COVID-19 myocarditis cardiac autopsy tissue for C4d (left, brown) and CD68 (right, brown). Corresponding areas from serial sections are shown.
